# Behavioural separation of face memory and face perception

**DOI:** 10.1101/2025.02.17.638018

**Authors:** Jan Kadlec, Catherine R. Walsh, Meytal Wilf, Jesse Rissman, Michal Ramot

## Abstract

A long-standing debate in neuropsychology concerns whether perception and memory function as independent systems or interact to support cognition. To investigate this, we developed the Face Memory and Perception (FMP) task, a novel paradigm designed to systematically disentangle whether and how these processes interact under different conditions. Across five independent datasets with over 900 participants in total, we observed consistent evidence that face perception and working memory operate independently when task demands are low, but in more complex conditions, these processes appear to interact. Notably, this interaction emerged only when the interference directly involved face-processing mechanisms, and did not arise from a general increase in cognitive load. Rather than the use of shared resources by overlapping cognitive processes, this interaction was driven by a shift in behavioural strategy from holistic to feature-based face processing as a result of maintenance-disrupting interference. These results underscore the fundamental independence of perception and working memory while also explaining some of the conditions under which interactions might be observed.

## Introduction

The traditional modular view of cognitive functions posits that perception and memory are distinct cognitive functions, each supported by specialised brain regions: the ventral visual stream is dedicated to visual perception, while the medial temporal lobe, including the hippocampus, supports declarative memory^1–5^. This view is supported by cases such as acquired prosopagnosia or apperceptive agnosia, where disruptions in memory or perception occur selectively without necessarily affecting the other, providing evidence for the dissociation of these processes at the neural level^6–10^. However, this strict separation has recently been questioned (see reviews^11–14)^. Mounting evidence suggests that the neural circuits and processes for perception and memory are not completely distinct and that they overlap and interact continuously. Thus, the more pertinent question might not be whether they are independent but rather which specific aspects of perception and memory are actually distinct.

To explore this issue, we turn to face memory and face perception, two processes central to our experiences in daily life. In the real world, these processes are inherently linked and most natural behaviours, such as face recognition, will involve elements of both. Judging traits such as gender, race and apparent age^15^ require not only the extraction of physical characteristics, but also reference to previous knowledge and templates, likely involving memory processes. This interaction between perception and memory is even clearer in the case of more nuanced social cues^16^ such as facial expression, mood, trustworthiness^17^, competence^18^, and intelligence which are very quickly extracted from complex face stimuli, along with elaborate first impressions^19–22^. One of the defining goals of the face system, recognising a familiar face, is perhaps the greatest example of the degree to which perception and memory processes are intertwined. Similarly in the brain, the distinction between these two processes is unclear. Large-scale networks throughout the brain, as well as specific brain regions, are specialised for faces or have been shown to be involved in face processing^23,24^, spanning the fusiform and occipital face areas^25,26^, superior temporal sulcus^27^, anterior inferior temporal area^28,29^, medial temporal lobe^30^ as well as the hippocampus, amygdala and entorhinal cortex. How, or whether, these networks are specialised for either perception or memory though, remains unclear.

Currently, there is no clear answer in the literature as to whether face perception and face memory are distinct, independent processes. On the one hand, some studies have shown that the two processes are largely correlated and appear to involve overlapping brain networks^31^, making them almost impossible to separate. We have also previously shown that individual differences in performance on face recognition tasks are predicted by networks encompassing regions involved in both memory and perception^24^. Nevertheless, studies across the lifespan show that face perception develops earlier than face memory^32^ as memory systems develop gradually^33^. Other studies show a differential effect of age on face memory and face perception^34^. Even stronger evidence for separation between face perception and memory comes from patients suffering from developmental or acquired prosopagnosia^35–40^, a condition involving a selective impairment in face recognition (see reviews^41,42^). Similar to general object perception and memory, some research in faces also separated prosopagnosia into *apperceptive*^8,43^ (patients cannot form an accurate representation of the face), *amnestic*^44,45^ (face perception is relatively accurate, but facial memory is impaired or lost) and *associative*^43^ (problem linking perception to memory; see reviews^9,35,37,46^ for details). Along similar lines, patients with isolated hippocampal damage often retain the ability to process and distinguish between unfamiliar faces or match them across short delays, suggesting that basic face perception and immediate matching rely on regions outside the hippocampus^47–49^. Although this separation has lately been criticised^41,50,51^, there seems to be a level of separation indicating dissociable neural correlates^52–54^ (see review^55^). Finally, there have also been more recent attempts to show the separation of face memory and face perception in healthy populations. These studies either regress variance from face perception^56^ obtained on purely perceptual tasks or use additive-factor logic to claim independence^57^. While both studies agree that face perception and face memory are distinct cognitive processes, Stantić suggests that, while separable, the two abilities may overlap in certain populations, particularly those with extreme face recognition abilities. In contrast, Hacker’s study emphasises that this separation is robust even in typical populations, highlighting independence as a likely universal cognitive principle across face and object recognition.

In general, the literature suggests that the dissociation between perception and memory is not clear-cut and that they can, but do not necessarily, interact. We therefore set out to systematically study both processes in a unified task framework in order to elucidate the conditions that give rise to interaction. To do this, we developed a new Face Memory and Perception (FMP) task, based on a match-to-sample paradigm, that independently modulates face memory difficulty (focusing on working memory) and face perception difficulty (through morphing). The FMP allows us to separately examine working memory demands and perceptual difficulty, as well as factors like cognitive load and interference. The basic premise of the FMP design is that if face perception and memory rely on shared cognitive resources, increasing perceptual difficulty should compound the effects of memory difficulty, leading to a significant interaction. Conversely, if these processes operate independently, their effects should be additive rather than interactive. Thus, the presence or absence of an interaction between perceptual and memory difficulty provides key evidence regarding the degree of independence between face perception and face memory. We collected five independent datasets, with 907 participants in total. Across these datasets, we systematically varied key experimental parameters—such as stimulus properties, interference conditions, and randomisation schemes—to confirm that our effects were not dependent on specific task settings. Despite these variations, we found that face memory and face perception are consistently separable across all datasets under most conditions but interact when there is interference that directly affects face processing. Surprisingly, this interaction was not driven by cognitive load per se or by the visual complexity of the distractor stimuli, but rather stemmed from participants switching between different holistic versus feature-based face processing strategies. Our results reinforce the basic independence of memory and perception and provide insight on the origin of their interaction under more difficult conditions, suggesting that it stems not from an overlap in cognitive resources, but rather from a behavioural shift. Understanding the basic mechanisms underlying these processes is crucial for better characterising and treating deficits in memory and perception across all domains.

## Results

### Independence of face memory and face perception in low-demand conditions

The experimental design of the FMP task is tailored to evaluate the interaction between two key variables: memory load and perceptual difficulty. We employed a 3x3 task design, modulating both perceptual difficulty and memory load independently. In all conditions, participants are first presented with a cue stimulus, followed by two probe images that appear below the cue (creating a triangle layout). One probe image is an exact match, and the other is a morphed version of the target image. We manipulated perceptual difficulty (Fig. 1a) by morphing faces to create three levels of difficulty: 24% morph (most difficult), 30% and 36% (easiest; see Methods for further details on the morphing procedure). Orthogonally, memory difficulty was modulated by changing the working memory demands during the task (Fig. 1b). In the easiest Matching (M) condition, the cue stimulus remains visible on the screen throughout the trial together with the two probes, and participants simply match the cue face with one of the two options. The more difficult Unfilled Delay (UD) condition adds a 4-second blank screen after the presentation of the cue stimulus, following which only the two probes appear, and participants have to select the correct face from memory. The third and most challenging condition, Filled Delay with emotional interference (FD-e), introduces an emotional face as a distractor during the delay. Participants are required to identify and remember the type of emotion portrayed in the image, thus increasing memory load and adding an additional cognitive demand. Following identification of the cue stimulus (as in the UD condition), participants are presented with two words describing emotions (e.g., happy sad), and must choose the label best describing the emotion presented in the distractor image (Fig. 1b in red).

**Figure 1.**
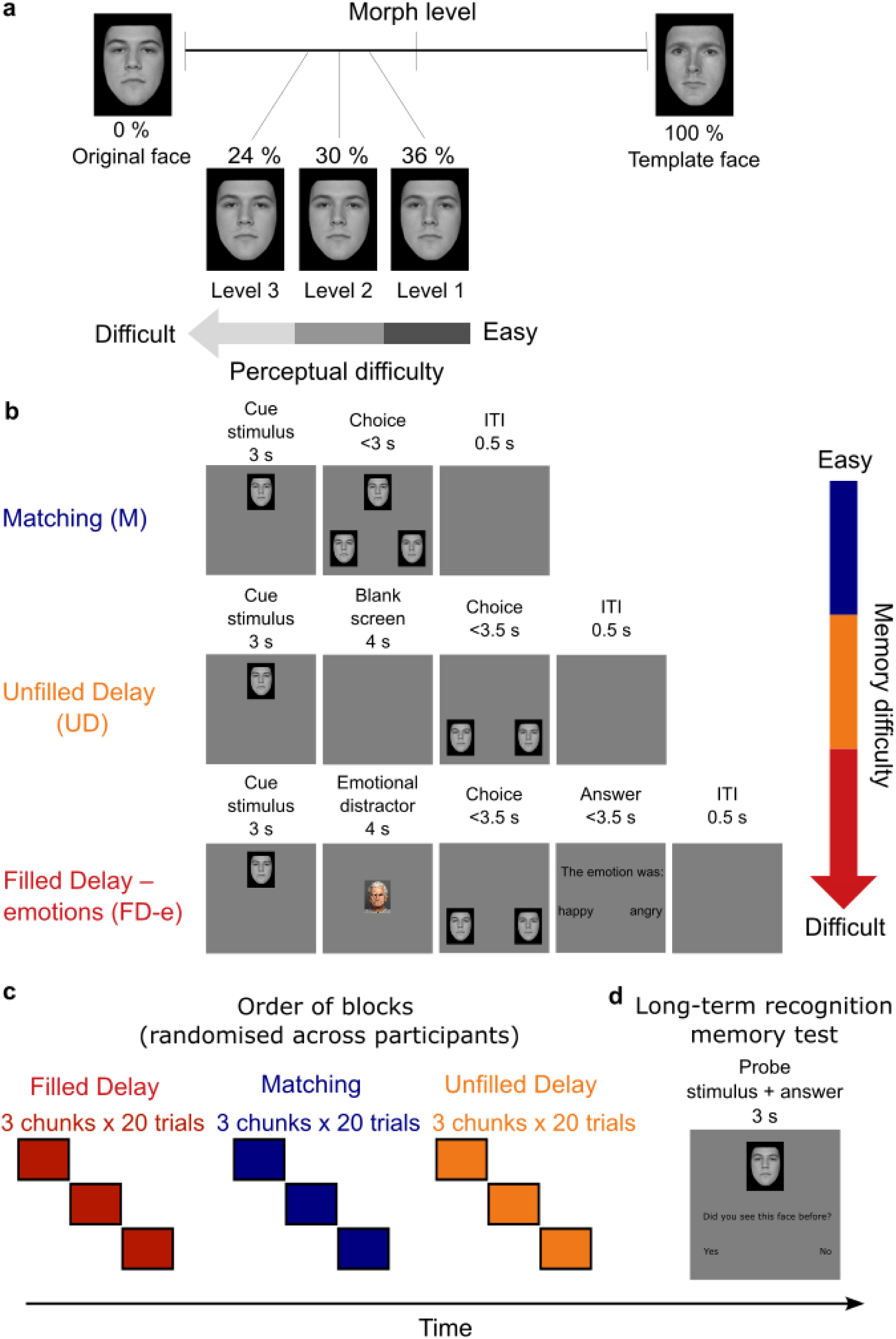
FMP Experimental design. **a.** The FMP parametrically modulates perceptual difficulty using face morphs of real face images. Perceptual difficulty is controlled by changing the morph level of the face. The greater the degree of morphing, the easier it is to distinguish between the two probes. The FMP uses three levels – 24% morph (hardest, level 3), 30% morph (level 2), and 36% morph (easiest, level 1). All male and female faces are morphed towards the same male and female template face, respectively. **b.** The FMP task modulates memory difficulty using different working memory demands. In blue, the easiest memory level, Matching (M), requires minimal to no memory. At first, a cue stimulus appears on the screen for 3s. Following that, two stimuli, one of which is an exact match and the other is a slightly morphed version of the face, appear underneath, forming a triangular arrangement. Participants are asked to choose which face matches the target face. In orange, the second memory condition introduces a delay of 4s after the initial presentation of the cue stimulus, during which a blank screen is shown (Unfilled Delay, UD). After this delay, the two probes appear as they would in M, but no cue stimulus is present, and participants are asked to recall from memory which of the probes matches the cue stimulus. In red, the hardest memory condition introduces active interference with the face maintained in memory by presenting another face during the 4s delay. This face is different from the cue and probes in that it is in colour, includes hair, and bears a clear emotion (Filled Delay – emotions, FD-e). Participants are instructed to remember the emotion displayed by the distractor face, as they will be asked about it later. **c.** The FMP experiment employs three levels of perception fully crossed with three levels of memory design. Each perceptual level x condition contains 20 trials, yielding a total of 180 trials. The order of conditions is randomised across participants. **d.** The long-term recognition memory test is performed at the end of the experiment. Participants are presented with 60 faces (30 new, 30 old, drawn from the first memory condition encountered). Images used in this figure are used in accordance with copyright and were part of the London^58^ and the FACES^59^ databases.

The factorial design fully crosses three memory conditions with three levels of perceptual difficulty, creating 180 trials in total (Fig. 1c). Each memory condition is divided into three chunks to mitigate fatigue, with images from all three perceptual difficulty levels randomly distributed across all three chunks. Images never repeat across trials and the order of memory conditions is randomised across participants. Participants always complete all 60 trilas of a condition before moving on to the next. Additionally, a final long-term recognition memory test assesses participants’ recognition of 60 faces, half previously seen and half new, gauging both recognition accuracy and confidence (Fig. 1d). This design allows us to examine not only whether perception and memory interact generally, but also how and if such interaction might differ as a function of cognitive load. While we expect to see a main effect of both perception and memory load (decreasing accuracy as the task becomes more difficult), it is not clear whether there will be an interaction between these two factors and whether this interaction will be evident across all conditions. The final “old”/”new” recognition task allows us to link long-term memory to performance on these conditions and better understand the encoding that takes place in the different memory conditions.

To control for different image properties and experimental design decisions, we collected three independent datasets (see Methods). First, we gathered data from N=259 participants (Dataset 1, Fig. 2a). To account for the slightly divergent effect of the morphing procedure on different images, which also resulted in differential charges in image entropy (a measure of the local variation among neighbouring pixels, see Methods for a detailed description and examples in Supplementary Fig. 1), we created a fully randomised stimuli set with entropy-matched stimuli, involving N=153 participants (Dataset 2, Fig. 2b). Finally, we investigated the impact of cue stimulus identity (whether the target stimulus is the unmorphed or morphed image), discussed in the next section, with data from N=262 participants (Dataset 3, Fig. 2c). Figure 2 displays the results from each of the three datasets separately, as well as the overall results from combining all three independent datasets (Fig. 2d). As expected, we observed main effects of perception and memory load in each dataset separately and in the concatenated dataset (2-way ANOVA significant main effect of memory condition: F(2, 1332) = 894.83, p < 0.001; 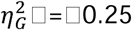; significant main effect of perceptual difficulty: F(2, 1332) = 474.11, p < 0.001; 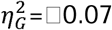 for the concatenated dataset, see SI Tables 1-6 for ANOVAs of all datasets). If face memory proficiency depended on face perception proficiency (or vice versa), an increase in memory difficulty would influence the difficulty of the perceptual task, leading to a statistical interaction between the two. We tested this hypothesis using the three independent datasets (Fig. 2a-c) as well as a concatenated dataset of all three datasets (Fig. 2d), and indeed found a significant interaction of memory condition and perceptual difficulty in all of the datasets. Specifically for the concatenated dataset: F(4, 2664) = 33.01, p < 0.001; 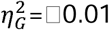 and see SI Tables 1-6 for all datasets.

**Figure 2.**
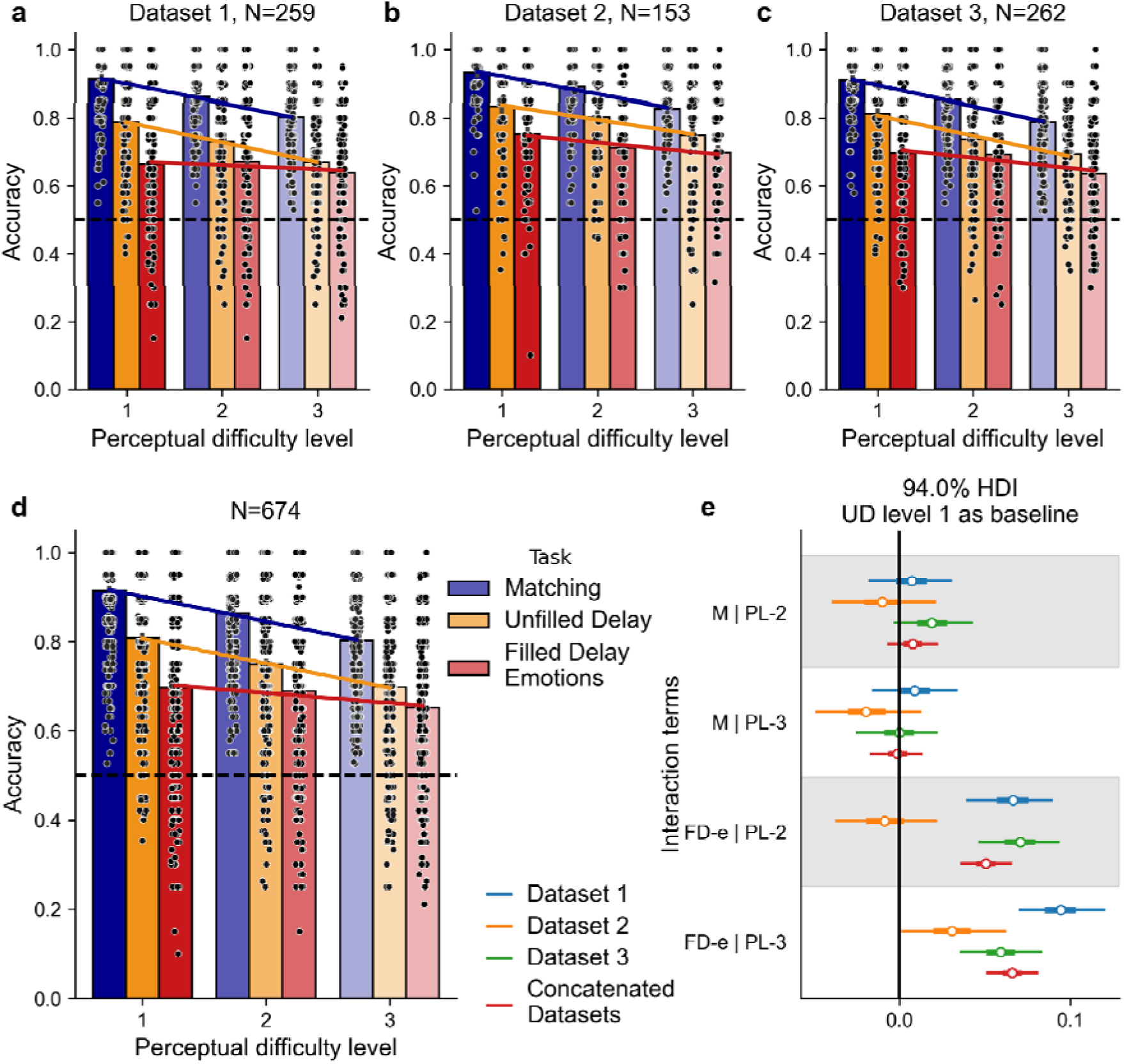
FMP results show independence of Matching (M) and Unfilled Delay (UD) across perceptual difficulty levels (PLs) but not of the Filled Delay (FD-e) memory condition and perceptual difficulty. Each dot represents a single participant’s score that is computed as the mean of 20 trials. As in Figure 1, perceptual difficulty levels are as follows: level 1 (PL-1, easiest, 36% morph), level 2 (PL-2, 30% morph), and level 3 (PL-3, hardest, 24% morph). **a-c** three independent datasets, see Methods for details. **d.** Concatenated datasets 1, 2, and 3. Connecting lines serve as a visualisation of interactions, where parallel lines reflect independence of memory and perception (M and UD), while non-parallel slopes across memory conditions reflect interaction between the processes (FD-e and M, FD-e and UD). **e.** Posterior distributions of interaction terms across the three datasets and the concatenated dataset show no significant interaction between UD level 1 and M at either of the perceptual difficulty levels 2 or 3 (M | PL-2 and M | PL-3, respectively; posterior distributions contain 0 in their 94% HDI) and that there is a significant interaction between UD and FD-e and perceptual difficulty except for Dataset 2 on level 2. UD perceptual difficulty level 1 is set as baseline.

It is difficult to pinpoint the source of the interaction using frequentist statistics such as ANOVAs when there are many potential interacting levels such as in our fully crossed 3x3 design. We therefore used Bayesian modelling to investigate the nature of the interaction between performance on different memory load conditions and different perceptual difficulty levels. First, we set the UD and perceptual difficulty level 1 (PL-1) as the reference level and validated the ANOVA results by showing that the Bayesian models revealed significant main effects for both memory conditions and perceptual difficulty, as indicated by the 94% high-density interval (HDI) of the posterior distributions not including zero (Supplementary Fig. 2a-d). The HDI is the Bayesian analogue to the frequentist confidence interval and it means there is a 94% probability that the true parameter value falls within the specified range, indicating a meaningful and reliable effect. We considered a fixed effect or interaction significant if its 94% HDI did not include zero, as done recently^60,61^. Subsequently, we compared a model that included an interaction term to a model without an interaction term (see Methods for details). If the interaction played a significant role, we would expect the model with the interaction term to perform better and explain more variance than the model without this term. This is what we observed in three out of the four tested datasets, where the model incorporating the interaction term provided a better fit compared to the model without the interaction (Supplementary Fig. 3a,c,d, Methods). In Dataset 2 (Supplementary Fig. 3b), this difference was not meaningful.

To identify the origin of the interaction, we examined the posterior distributions from the interaction model, which revealed a significant interaction for FD-e vs. UD across all three datasets, as well as the concatenated dataset (Fig. 2e). We did not observe a significant deviation from zero for M vs UD, suggesting that face perception and memory operate independently in the absence of additional cognitive demands. In contrast, the significant interaction observed for FD-e vs UD suggests that these processes can influence one another when participants are required to process and retain a face distractor’s emotion. Furthermore, the posterior distributions also clarified the non-significant difference observed in Dataset 2. While there was a significant interaction between FD-e and UD at perceptual level 3, there was no significant interaction between FD-e and UD at perceptual level 2. This diminished effect of the interaction caused the non-significant difference between the two models (Fig. 2e; see Methods for details).

The above analysis gave us all interactions between UD and the other memory conditions. For completeness, we also compared all other conditions to the M condition. We observed the same pattern across all datasets, mirroring the results of the previous analysis: there was a significant main effect of memory and perceptual difficulty (Supplementary Fig. 2e-h), no interaction between UD and M, and there was a significant interaction between FD-e and M on all levels in all datasets, except for FD-e level 2 in Dataset 2 where we find no interaction (Supplementary Fig. 4).

To validate that the observed interaction in the FD-e condition does not stem from a floor effect or restricted range since this is the most difficult condition, we selected the top 50% of participants on the FD-e condition and reran the analysis. Performance on FD-e level 1 in this subsample of participants (0.80 ± 0.10, proportion correct ± standard deviation) was comparable to that of the full sample on UD level 1 (0.81 ± 0.14). The results, shown in Supplementary Figure 5, corroborate results from the full sample – independence between M and UD on all levels and interaction between UD and FD-e on all levels and in all datasets except for Dataset 2 and perceptual level 2 (as before).

Even though the order of the conditions was counterbalanced across participants, we wanted to test if there were any learning or exposure effects, and if these might explain any part of our results. We therefore repeated the main analyses on three subsets of participants: those who began with the M condition, the UD condition, or the FD-e condition, respectively. As shown in Supplementary Figure 6a–c, the results remain consistent across these subsets. Model comparisons and interaction patterns are preserved (Supplementary Fig. 6d, e), with no qualitative changes. We further examined whether condition order influenced performance by comparing accuracy across participants who completed the M/UD/FD-e conditions first, second, or last. While a significant order effect was found for the M condition (F(2, 671) = 3.20, p = 0.041; 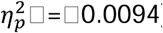), post-hoc analysis revealed this was driven by slightly higher accuracy when the condition appeared first compared to second (Post-hoc Tukey test: first vs second, T=2.35, p=0.049, Cohen’s d=0.23), but not compared to last (Post-hoc Tukey test: first vs last, T=1.96, p=0.12, Cohen’s d=0.19; last vs second, T=0.42, p=0.90, Cohen’s d=0.04, see SI Table 7). Note that this comparison is across different participants, and most likely reflects either minor differences between the groups, or a possible small effect of fatigue. In either case, the result is inconsistent with a learning effect (which would predict increased performance with later condition order), accumulation of interference (which would predict decreased performance with later condition order), see Supplementary Figure 6f.

### Lack of perceptual symmetry between morphed and unmorphed cue stimuli

Our results thus far suggest that perception and memory processes are fundamentally independent, yet they interact when interference momentarily disrupts working memory maintenance. These results are remarkably consistent across all three datasets, despite the fact that we changed different image properties and image randomisation schemes. One aspect of image manipulation that we did not test relates to the symmetry of the morphing. From a theoretical perspective, presenting the unmorphed image or the morphed image as the cue stimulus should be entirely symmetrical. Mathematically, the visual distance between the unmorphed and morphed probes is identical in both cases, meaning accuracy should not differ. However, it is possible that the perceptual distance is not truly symmetrical, and changes in perceptual difficulty could influence the nature of the interaction. We therefore took a closer look at Dataset 3 with N = 262 participants, in which we manipulated whether the cue stimulus was the unmorphed image, or the morphed image. The unmorphed image was presented as the cue stimulus in 73% of trials, while in the remaining 27% the morphed image was presented (Fig. 3a). Participants were not informed about this manipulation, and their instructions remained the same, i.e., to choose an exact match.

**Figure 3.**
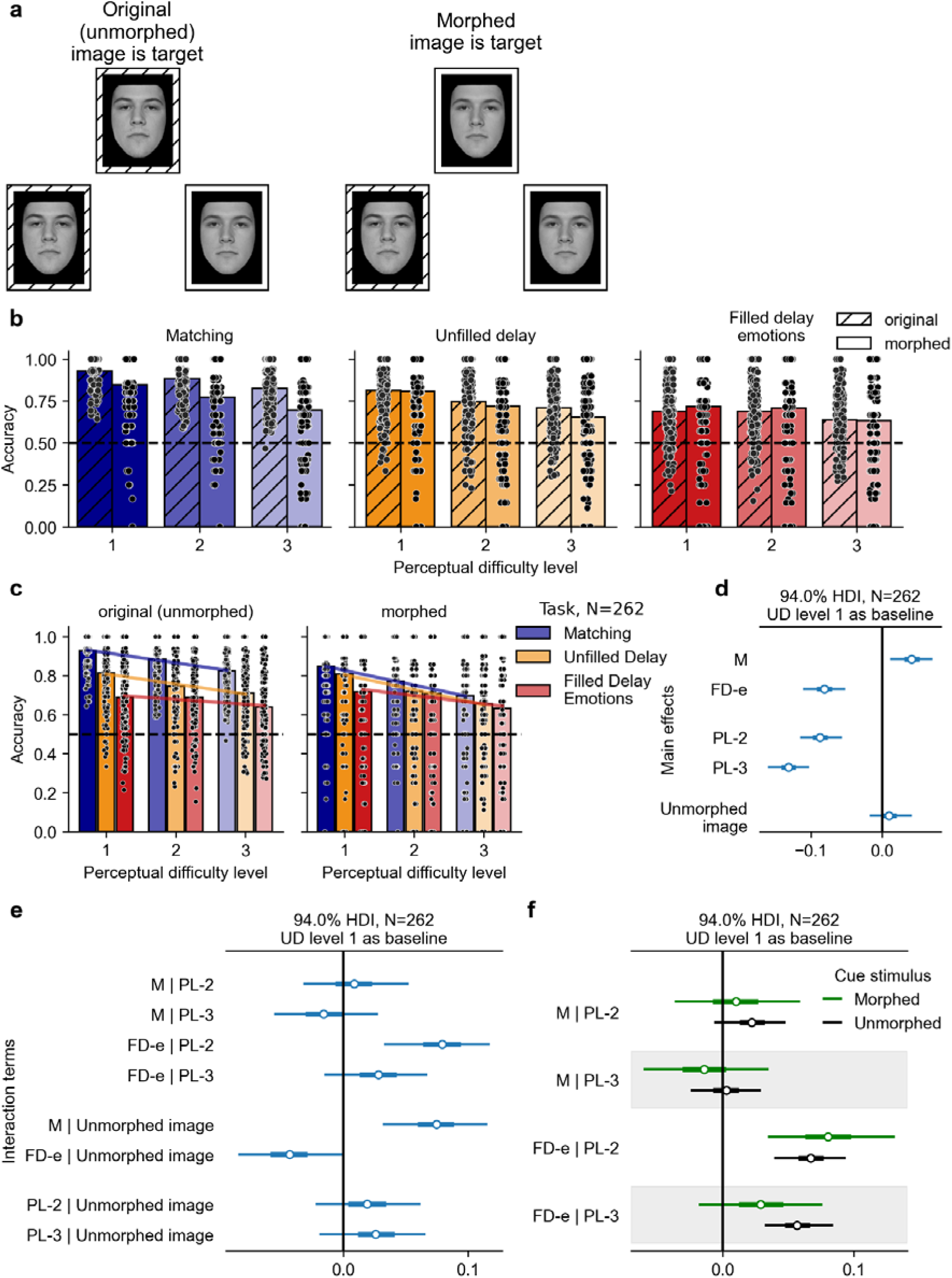
Morphed and unmorphed cue stimuli differentially affect accuracy on but not interaction between memory conditions and perceptual difficulties. **a.** Experimental design. In all memory conditions, participants need to find an exact match to the cue stimulus presented initially. In 70-76% of trials, this cue stimulus is the original (unmorphed) face as in previous datasets (left). In contrast, in the rest of the trials, the morphed face is used as a cue stimulus (right). Images used in this figure are used in accordance with copyright and are part of the London^58^ database. **b-c.** Accuracy across memory and perceptual difficulty in Dataset 3 (N=262) split by cue stimulus grouped either by memory condition (**b**) or cue stimulus morphing (**c**) shows a significant effect of cue stimulus morphing in M but not in UD, and in FD-e, albeit in the opposite direction. Despite the impact on accuracy, the cue stimulus doesn’t change the independence of M and UD, as there is no interaction between M, UD and the perceptual difficulty levels (panel **e**). **d.** There is no main effect of cue stimulus in the UD condition (Unmorphed image contains zero in its 94% HDI, the UD perceptual level 1 with morphed cue stimulus is set as a reference level) and as before, there is a significant effect of both memory (M, FD-e) and perceptual (PL-2, PL-3) difficulty levels. **e.** The results corroborate no interaction between M and UD (at both perceptual levels, M | PL-2 and M | PL-3), significant interaction between UD and FD-e (only at perceptual level 2, FD-e | PL-2) and further shows a differential effect of cue stimulus (Unmorphed image) on face memory (M | Unmorphed image and FD-e | Unmorphed image) but not face perception (on both perceptual levels, PL-2 | Unmorphed image and PL-3 | Unmorphed image contain 0 in their 94% HDI). This interaction between the cue stimulus and the memory levels goes opposite directions for M and FD-e. **f.** Posterior distributions of interaction terms show no significant interaction as the 94% HDI contains zero in both subsets of Dataset 3 between UD and FM with UD set as reference level. The FD-e condition behaves differently in the dataset with morphed cue stimuli as there is no significant interaction between UD PL-1 and FD-e at perceptual level 3 but there is at perceptual level 2. Interactions between FD-e and both UD and M are significant for PL-2 for both types of cue stimuli. Interactions at perceptual level 3 are significant between FD-e and both M and UD only when the unmorphed stimuli is the target (Supplementary Fig. 8c).

We found that accuracy changed across conditions depending on whether the morphed stimulus is used as the target, yet the interactions between memory and perception were not altered. Figure 3b-c shows the same data from Dataset 3 as shown previously in Figure 2c but now split by condition (left, unmorphed image as target; right, morphed image as target). There was a significant effect of cue stimulus in the M condition (Fig. 3e where 94% HDI interaction between M and unmorphed image doesn’t contain zero and Supplementary Fig. 7 as the main effect when M is set as a reference level) but not in UD (Fig. 3d). Interestingly, in FD-e, there was a smaller but still significant effect, though the direction of the effect was opposite to that found in the M condition, with accuracy being higher on trials where cue stimuli were morphed faces (Fig. 3e and Supplementary Fig. 7, 94% HDI interaction between FD-e and unmorphed image doesn’t contain zero).

Although the cue stimulus significantly and differently affects overall accuracy across various memory conditions, the independence of M and UD remains unaltered. This is evidenced in the lack of interactions between M, UD, and perceptual difficulty both when averaging across morphed and unmorphed cue conditions (see Fig. 2c) and when analysing each condition separately (see Fig. 3c,f and Supplementary Fig. 8c). As in Dataset 1 (and in this dataset averaged across morphing conditions), there was a significant interaction in FD-e with difficulty level 2, when comparing to both UD and M (Fig. 3e and Supplementary Fig. 7, 94% HDI does not include zero). Further investigation into the effect of the cue stimulus showed that only when the cue stimulus was morphed did we observe no interaction between FD-e and difficulty level 3, perhaps because of the lower number of trials available in this condition. This further replication of the fundamental finding – despite the interaction caused by whether the cue stimulus was morphed or not – that memory and perception are independent under the M and UD conditions, but not under the FD-e condition, provides additional evidence regarding the crucial role of task demands in driving the interaction.

### Cognitive load, per se, does not drive the interaction

A simple explanation of our results thus far is that perception and memory interact as cognitive load increases. We therefore set out to determine whether the nature of the cognitive load is important in generating this interaction. The FD-e condition differed from the UD condition in two key aspects – first, there was an additional non-memory cognitive demand which directly involved face processing (recognise the emotion of the face), and second, there was increased memory load (remember the emotion as well as remember the original cue stimulus). We tested whether engaging face processing directly in the higher load conditions is necessary, or whether simply increasing cognitive load and memory demands in an orthogonal manner will have similar effects. To do this, we introduced a new condition, Filled Delay with Math distractor (FD-m) that had similar demands to FD-e but did not involve face processing (Fig. 4a). In FD-m, participants were required to evaluate a simple mathematical equation (non-memory cognitive demand) during the delay period, and remember whether it was true or false (increased memory load). We hypothesised that the FD-m condition would show poorer performance than the UD condition but would not disrupt face memory as much as the FD-e condition. Contrary to expectations, our results indicated that performance in the FD-m condition was indistinguishable from the UD condition (Fig. 4b). This finding was further corroborated by Bayesian modelling, which shows no difference between these conditions (Supplementary Fig. 9c).

**Figure 4.**
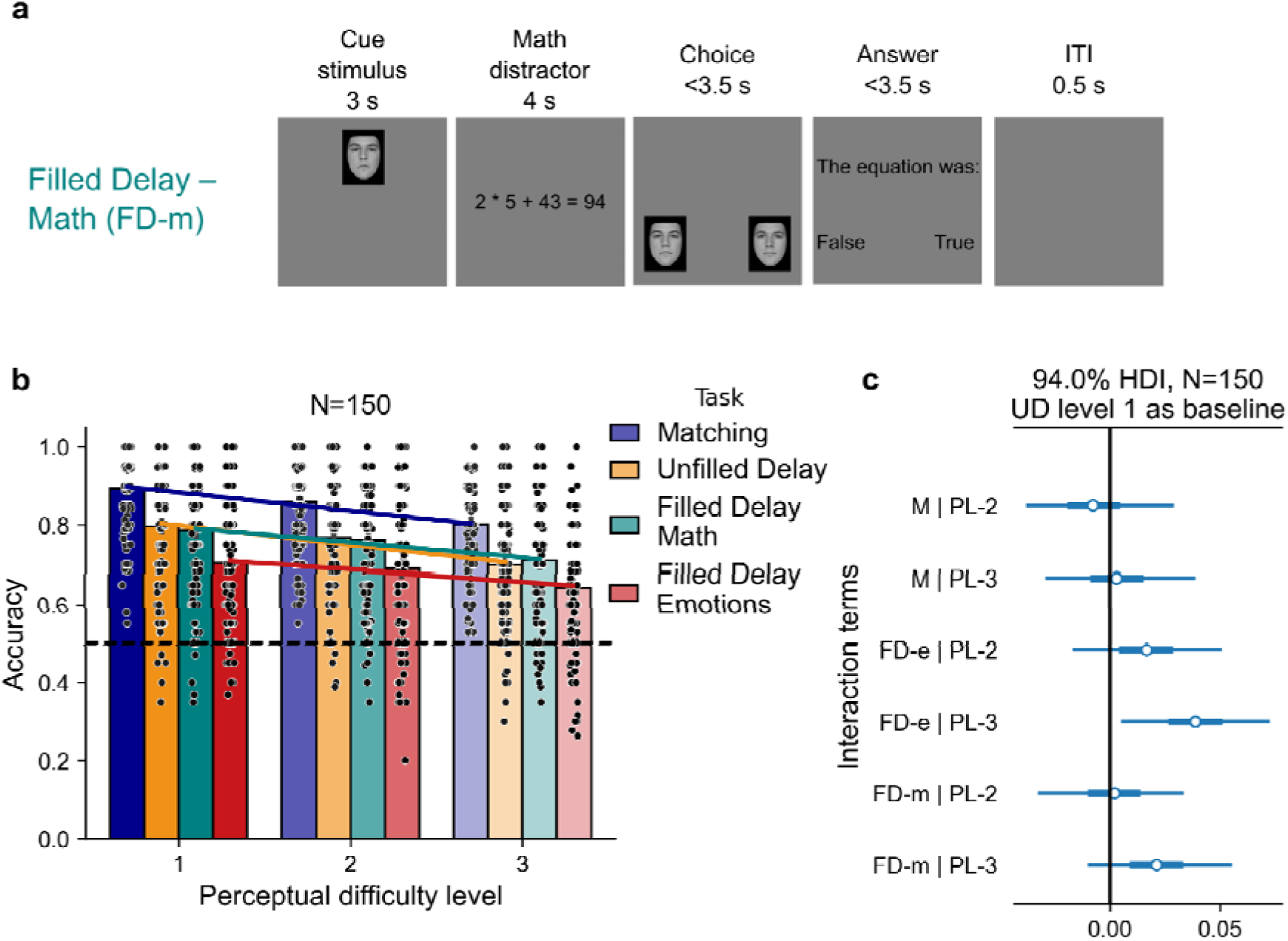
Orthogonal filled delay with math distractor (FD-m) is not different from Unfilled Delay (UD). **a**. Experimental design. In teal, this memory condition introduces an active interference that is orthogonal to face processing (FD-m). During the 4s delay that follows the initial face presentation, participants view a simple arithmetic equation and are asked to remember whether this equation was True or False. Images used in this figure are used in accordance with copyright and were part of the London^58^ databases. **b.** Results of Dataset 4 show the same accuracy on UD and FD-m memory conditions. Furthermore, both UD and FD-m show no interaction with M, corroborating results from Figure 2 and the previous datasets. As in previous datasets, the FD-e condition also had a significant interaction with perceptual difficulty in both M and UD on level 3. **c.** Posterior distributions of interaction terms show no significant interaction as the 94% HDI of all of them contain zero (black line) except for FD-e level 3. Interaction terms are defined as condition:difficulty level. Zero corresponds to UD level 1.

There is a small but significant difference between models with and without an interaction term in Bayesian modelling (Supplementary Fig. 9a). Looking at the posterior distributions, we found that the only significant interaction was between UD and FD-e at perceptual level 3 (Fig. 4c, Supplementary Fig. 9b). We did not observe any interactions for the FD-m condition (Fig. 4c). These results are consistent with the findings across all datasets (Fig. 2), underscoring the independence of M from UD and suggesting that interactions do not generally manifest when memory constraints and perceptual difficulty are high but rather also require higher cognitive load specifically involving face processing. The previously observed significant interaction between the FD-e condition, and perceptual difficulty in the UD and M conditions was observed in this analysis as well (Fig. 4c, Supplementary Fig. 9b). Similarly to before, no significant interaction was observed when we set the M condition as a baseline except for an interaction of FD-e and M on perceptual level 3, mirroring the interaction of FD-e and UD on perceptual level 3.

### Non-specific visual interference does not drive the interaction

Another possible explanation could be that it is the visual interference – processing highly complex visual stimuli (emotional faces), rather than the cognitive nature of the task, that produces the interaction. The FD-m condition lacks visual complexity, and therefore cannot address this possibility. To test this hypothesis more directly, we introduced a new condition, FD-v (Filled Delay with Visual non-face distractor), in which participants viewed images of food plates during the delay period and had to classify whether the food was vegetarian or not, a question they were asked following the probes, as in the FD-e and FD-m conditions (Fig. 5a). Food plates were selected as visually complex and socially salient stimuli, approximately matched to faces in real-world size and perceptual richness. The stimuli were piloted to ensure that categorisation difficulty was comparable to the emotional judgement task used in the FD-e condition (see Methods).

**Figure 5.**
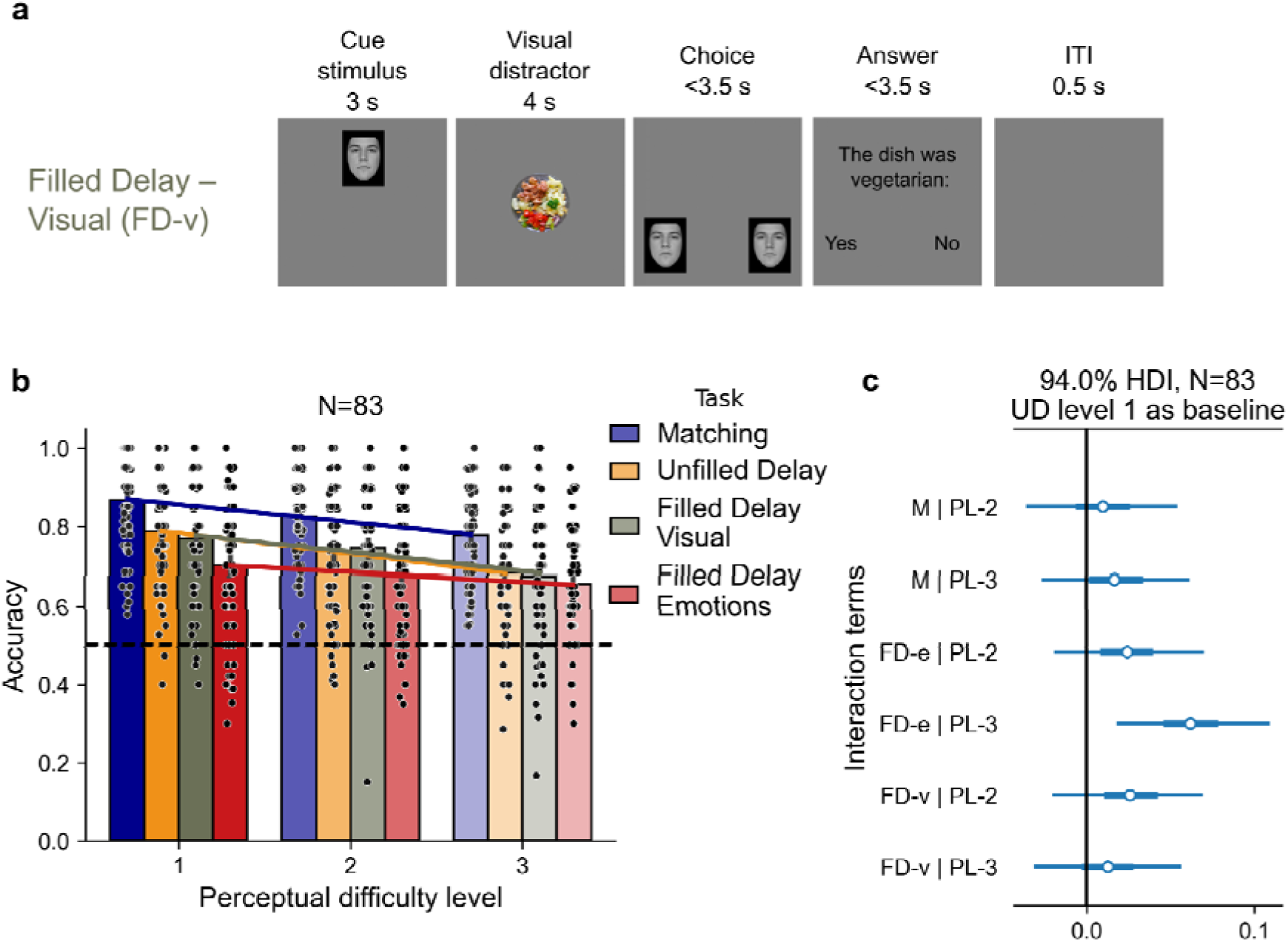
Orthogonal filled delay with rich visual (non-face) distractor stimuli (FD-v) is not different from Unfilled Delay (UD). **a**. Experimental design. In olive, this memory condition introduces an active interference that is orthogonal to face processing (FD-v). During the 4s delay that follows the initial face presentation, participants view a food plate and are asked to decide and remember whether this food was vegetarian (Yes) or not (No). Images used in this figure are used in accordance with copyright and were part of the London^58^ databases or taken by JK. **b.** Results of Dataset 5 show the same accuracy on UD and FD-v memory conditions. Furthermore, both UD and FD-v show no interaction with M, corroborating results from Figure 2 and Figure 4 and the previous datasets. As in previous datasets, the FD-e condition also had a significant interaction with perceptual difficulty in both M and UD on level 3. **c.** Posterior distributions of interaction terms show no significant interaction as the 94% HDI of all of them contain zero (black line) except for FD-e level 3. Interaction terms are defined as condition:difficulty level. Zero corresponds to UD level 1.

Behavioural results showed that performance in the FD-v condition was indistinguishable from that in the UD condition across all levels of perceptual difficulty (Fig. 5b), a finding further supported by Bayesian modelling, which revealed no significant difference between FD-v and UD (Supplementary Fig. 10). This pattern closely mirrors the findings from the previous section (Fig. 4) and applies to all aspects of the results described there. There is no difference between models with and without an interaction term in Bayesian modelling (Supplementary Fig. 10). Looking at the posterior distributions, we found that the only significant interaction was between UD and FD-e at perceptual level 3 (Fig. 5c, Supplementary Fig. 10b). We did not observe any interactions for the FD-v condition (Fig. 5c). The previously observed significant interaction between the FD-e condition, and perceptual difficulty in the UD and M conditions was observed in this analysis as well (Fig. 5c, Supplementary Fig. 10b). Similarly to before, no significant interaction was observed when we set the M condition as a baseline except for an interaction of FD-e and M on perceptual level 3, mirroring the interaction of FD-e and UD on perceptual level 3.

Together, these findings replicate the pattern observed with the FD-m condition and reinforce the conclusion that the interaction between face perception and memory depends on domain-relevant interference, rather than non-specific visual content or general cognitive load.

### Holistic vs feature-based processing as a driver for the interactions

While we found an interaction between memory and perception in the memory condition with the emotional face filled delay, FD-e, this interaction was not present in the orthogonal filled delay condition, FD-m, arguing against a simple increase in general cognitive load as the driver for this interaction. According to cognitive load theory^62,63^, increased cognitive load should result in greater interference (and lower performance) with increasing difficulty, predicting a steeper decrease in accuracy as perceptual difficulty increases^64^. Surprisingly, and consistently across all five datasets and the concatenated dataset (Fig. 2-4), the interaction observed in the FD-e condition results from a shallower decrease in accuracy over the different perceptual levels, such that the hardest perceptual difficulty level 3 appears less impacted by the filled delay manipulation than level 1, rather than vice versa.

An alternative explanation for the origin of the interaction might lie in a shift in strategies. As perceptual difficulty increases, the probe faces become more similar. This might prompt participants to shift from a more holistic face processing approach to a more feature-based one. Unlike in the M, UD, and FD-m conditions, the emotional face used in the FD-e condition interferes with the representation of the cue stimulus in working memory, and might therefore selectively impact holistic processing across all perceptual levels^65^. Accordingly, we would expect all perceptual levels in the FD-e condition to be primarily reliant on feature-based processing. Consequently, at the easier perceptual levels, there is a greater effect of the manipulation introduced by the FD-e causing a greater decrease in performance in FD-e compared to the other conditions. As perceptual difficulty increases, this interference is diminished as participants already rely on feature-based processing, causing a smaller drop in performance. This pattern would explain the shallower performance slopes observed in FD-e, which drives the observed interaction, as well as the lack of interaction in FD-m.

To determine whether participants use global (holistic) or local (feature-based) processing, we utilised the inversion effect which has previously been used to estimate the extent of holistic vs. feature-based processing^66–69^, and collected an additional dataset of 71 participants. The inversion effect is defined as the difference in accuracy on trials where the face is upright, vs. trials where the face is inverted. A greater inversion effect would indicate more holistic processing, whereas a shift to feature-based processing would result in a decrease in the inversion effect. To avoid potential confounds from the additional interference in the FD-e condition, we tested the inversion effect on the UD condition, across all three levels of perceptual difficulty. Each difficulty level consisted of 40 trials per inversion condition, resulting in a total of 120 upright and 120 inverted face presentations. The results, illustrated in Figure 6, demonstrated significant main effects of both inversion and perceptual difficulty on accuracy, as well as a significant interaction between these factors (2-way ANOVA, inversion: F(1, 420)=62.57; p=2.3·10^-14^; 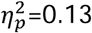, perceptual difficulty: F(2, 420)=20.26; p=4.1·10^-9^; 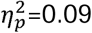, interaction: F(2, 420)=3.27; p=0.039; 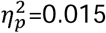). In line with the hypothesis that participants shift from more holistic to more feature-based processing as perceptual difficulty increases, the largest inversion effect is found for the easiest perceptual level, and the smallest inversion effect is found for the most difficult perceptual level, with post-hoc tests showing a significant difference between the easiest level and all other levels (Post-hoc Tukey test: level 1 vs 2, T=3.68, p=0.00077, Cohen’s d=0.41; level 1 vs 3, T=5.85, p=2.95·10^-8^, Cohen’s d=0.69; level 2 vs 3, T=2.17, p=0.077, Cohen’s d =0.28).

**Figure 6.**
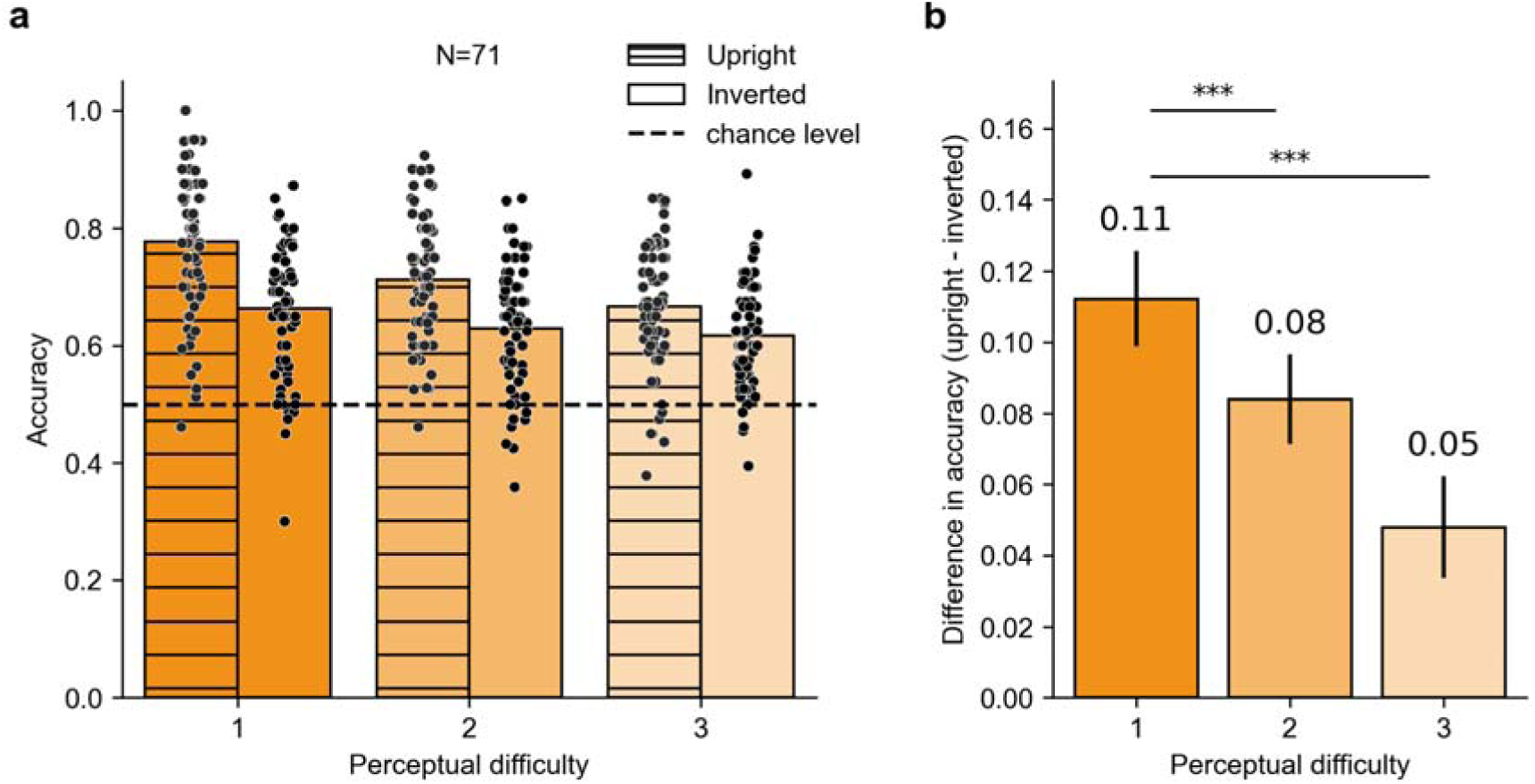
Differential inversion effect across perceptual difficulty levels. **a.** Accuracy in the UD condition across the three perceptual levels for N=71 participants. Lined bars show trials with upright faces and clean bars show trials with inverted faces. Dashed horizontal line denotes chance level. **b.** The inversion effect, defined as accuracy on upright minus inverted trials across perceptual difficulties. Error bars show standard error of the mean (SEM). Stars denote the significance of the difference between the different perceptual levels: *p*_<_0.001 (***).

### Long-term recognition memory

The conditions across our memory conditions, especially between the M and UD conditions, are overall very similar. This similarity raises the question of whether these conditions engage distinct underlying processes. As differences in encoding could impact the long-term retention of faces in memory, one potential measure for the distinction between the conditions is to assess the effects of the condition on long-term memory performance. As the M condition involves little to no working memory, and the FD-e condition actively interferes with the maintenance of working memory, we might expect that the encoding of the cue-stimulus in both the M and FD-e will be diminished compared to the UD condition, resulting in poorer performance on a long-term memory task involving those faces. To test this, we included a long-term recognition memory condition at the very end of the FMP (Fig. 1d). This task consists of 30 new, previously unseen faces and 30 old faces, drawn randomly from the first memory condition that the participant encountered. Importantly, the first memory condition is randomised across participants, so that for each participant, the “old” images were drawn from one of the three memory conditions. The time delay between first encountering the images, during the first memory condition which corresponds to the first block, and the old-new recognition task at the end of the experiment is approximately 30 minutes, allowing us to test the effects of encoding during the three conditions (M, UD, FD-e) on long-term memory performance.

This long-term memory test was added midway through data collection for Dataset 1, so only data from participants tested after its introduction were included in the analysis of that Dataset (Dataset 1b). As can be seen in Figure 7, consistently in all datasets but Dataset 2, d’ on images coming from both the M and FD-e conditions is significantly lower than on images from UD, suggesting that these conditions do indeed differ in how the stimuli are encoded (1-way ANOVA for concatenated Datasets: F(2, 593)=12.52; p=4.7·10^-6^; 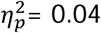, post-hoc Tukey test: FD-e vs M, T=0.73, p=0.75, Cohen’s d = 0.08, FD-e vs UD, T=-3.88, p=0.0003, Cohen’s d =-0.38, M vs UD, T=-4.66, p=1.2·10^-5^, Cohen’s d = -0.45, see SI Tables 8-11 for the remaining stats). Dataset 2 shows the same trend, though the differences do not reach significance, perhaps because Dataset 2 is the smallest of all datasets. This is a between-participant analysis, making it particularly sensitive to the relatively small sample size, with only 43 participants in the UD condition. Figure 7e shows the correlation between d’ in the long-term memory task and accuracy in each memory condition of the FMP on the concatenated dataset. Notably, this correlation is consistently stronger between UD and d’ than between d’ and either M or FD-e across all datasets separately as well in this concatenated dataset (Supplementary Fig. 11), though statistical tests for differences between correlation coefficients did not yield significance, likely due to their conservative nature.

**Figure 7.**
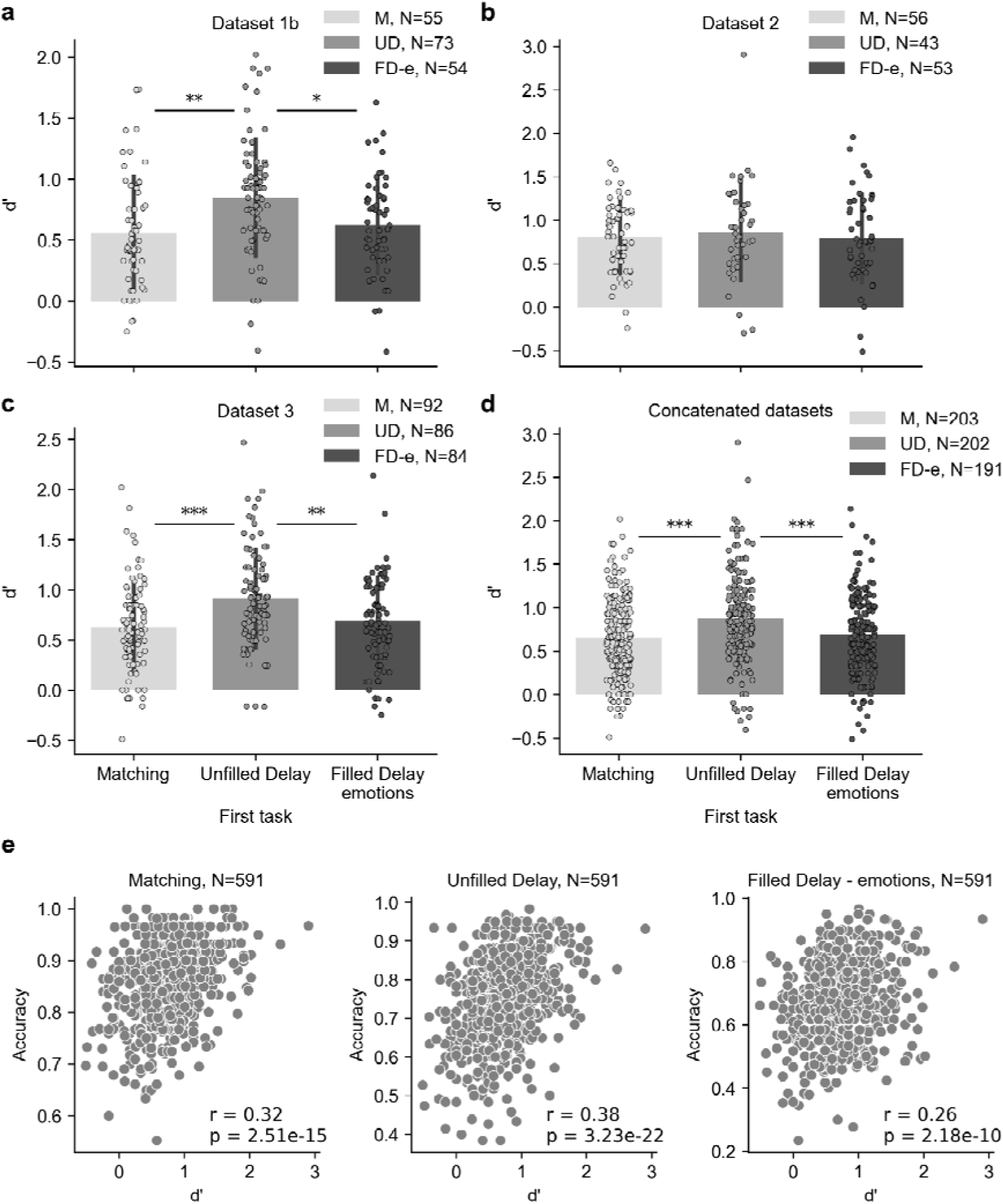
Long-term recognition memory shows differential encoding between the three memory conditions. d’ measure on the Long-term memory recognition task split by the memory condition from which the old stimuli were taken. Panels a-c correspond to the different datasets: Dataset 1b (for participants for which there was data collected on this task) (**a**), Dataset 2 (**b**), Dataset 3 (**c**), while panel **d** shows d’ when concatenating all three datasets together. Number of participants in each group is shown in the legend. The error bars represent standard deviation (SD), ** denotes p<0.01 and *** denotes p<0.001. **e.** Correlation between d’ and accuracy scores in the three memory conditions (M, UD, FD-e) in the concatenated dataset, regardless of which condition the “old” probes were drawn from.

## Discussion

Our findings provide valuable insights into the ongoing debate regarding the independence of memory and perception by systematically assessing their interaction under various conditions in one particular, highly studied domain – faces. Focusing on unfamiliar faces, we developed a large, diverse, standardised set of over 1,400 stimuli of real faces that reduced confounds like other-race effects, lighting inconsistencies, and prior exposure. Our new Face Memory and Perception (FMP) task (Fig. 1) leverages this exceptional stimulus set to parametrically study face recognition at different levels of perceptual difficulty and memory load, in order to pinpoint the conditions under which these processes interact.

Our large-scale data collection efforts, spanning five independent datasets comprising over 900 participants, not only allowed us to test multiple hypotheses, but importantly, it also allowed us to replicate our findings under different conditions. The first three datasets manipulated different image properties, while the final two datasets included a new distractor condition to test interference that is orthogonal to the face system (FD-m) and a control for the visual complexity of faces (FD-v). Despite these differences between the datasets, our main findings are remarkably consistent and were replicated across all five datasets, as well as in the concatenated dataset. In general, we found that working memory load does not interact with perceptual difficulty, as can be seen in the independence of the M (minimal/no working memory) and UD (working memory) conditions. Equally replicable across datasets was our finding that the FD-e condition (working memory plus active interference to the face system) does interact with perceptual difficulty (Figs. 2-5). This was true even in Dataset 3, where we observed large differences in performance depending on whether the morphed or unmorphed image was used as a cue. Although the cue stimulus effects differed between the memory conditions, there was no change in the overall interaction pattern between working memory load and perceptual difficulty, i.e., the effect of perceptual difficulty was independent of working memory load in the M and UD conditions, but not in the FD-e condition (Fig. 3). Across all datasets we therefore observe the same results – both the independence of perceptual and memory difficulty across the M and UD conditions, as well as an interaction when we introduce direct interference with face processing in the FD-e condition.

The fundamental independence of the M and UD conditions aligns with neuropsychological findings from prosopagnosia and prosopamnesia studies, which demonstrate dissociation between perception and memory impairments^45,52–55,70^. Our demonstration of behavioural independence, at least under specific conditions, strongly suggests that there exist distinct underlying neural mechanisms on which these processes rely, such as the ventral visual stream for visual perception^12^ and a distributed network for working memory^71–73^, for instance parietal regions and the prefrontal cortex^74–77^. Had the interaction between FD-e and perceptual difficulty been the result of increased cognitive load, even if only of specific load related to the face system, that would suggest that these same pathways may also be recruited by both processes as the task becomes more complex, or that additional shared processes are recruited. Our findings however point to a different mechanism for the interaction.

The FD-e condition, which involved interference during the delay period with an emotional face, introduced both additional non-memory cognitive demands, as well as increasing memory load. According to cognitive load theory^64^, easier tasks might rely on automatic processing that require less conscious attentional resources, while tasks with higher cognitive load will demand more allocation of attention and working memory. Cognitive load has been shown to have a complex effect on interactions in several cognitive domains, for instance in the cognitive-motor domain, higher cognitive load slows hand movements in a visuomotor task^78^ and reduces gait speed and stride length while performing a simultaneous dual-task^79^. This framework thus predicts a different relationship between perception and memory performance depending on the different levels of task cognitive loads, incurring more interactions as tasks become more difficult. Indeed, introducing emotional interference disrupted the distinct separation between face memory and perception in the FD-e condition, in support of the cognitive load theory.

On the other hand, under an orthogonal interference condition (FD-m) with similarly increased cognitive load, we did not find an interaction (Fig. 4). The absence of the interaction in the orthogonal interference condition still fits within the cognitive load theory, as according to the Working Memory Model^80^, the presence of an interaction might also be dependent on whether or not the additional task demands tap into the same or different processing subsystem as those of the basic task^81,82^. It is also possible that the affective nature of the task was the source of the interaction, as previous research indicates that emotional content can interfere with cognitive processing by engaging additional neural resources, such as the amygdala^83^, which is involved in both emotional processing^84,85^ and memory encoding^86,87^. These conceptual frameworks can account for the main differences between FD-e and FD-m conditions, as well as the lack of differentiation between the UD and FD-m conditions. However, they are less likely to account for the differences between the FD-e and the FD-v conditions (Fig. 5), as at least some of the visual processing mechanisms are shared, and food is likewise an affective stimulus. Even more compellingly perhaps, neither cognitive load theory nor the additional resources required for affective processing can explain the direction of the interaction that we observed, with the easiest perceptual levels being differentially more affected than the hardest perceptual level.

We therefore turned to search for alternative explanations for the origin of the interaction. The FD-v control, despite having similar task demands to and visual complexity as the FD-e condition, does not produce an interaction or a drop in performance and thus strongly suggests that active engagement of the face processing system drives the effect. It has previously been shown that the representation of faces in working memory relies primarily on holistic (gestalt) processing^88^, though recent works suggest that the type of task determines whether holistic or feature-based representations are used^89^. We hypothesised that increasing perceptual difficulty might cause participants to shift from a holistic processing strategy to using more feature-based information. The distractor in the FD-e condition interferes with face processing by specifically disrupting the holistic representation of the cue stimulus face^65,89,90^ that participants actively maintain in their memory. If easier perceptual levels are indeed more reliant on holistic processing, then they will be more affected by the emotional interference face, resulting in a larger decrease in performance on FD-e compared to the UD condition for the easier perceptual levels, and a smaller decrease compared with UD in the hardest perceptual level, in line with the interaction that we observed.

It is difficult to directly estimate the level of holistic processing vs. feature-based or configural processing (which was less relevant in this context), but a good proxy for this is the inversion effect. The inversion effect is the observation that inverting images causes a much greater drop in task accuracy for faces than for any other type of stimulus^91^ (see also reviews^66,69,92^). It has been argued that the inversion effect is evidence for the holistic processing of faces^93–98^. In particular, it has been posited that differences in the inversion effect across categories and conditions correspond to the level of holistic processing^99^. In our case, a pronounced inversion effect (i.e., a large difference in accuracy between upright and inverted faces) would indicate greater holistic face processing, whereas a smaller inversion effect would suggest greater reliance on feature-based processing. Our data in Figure 6, from a sixth independent dataset collected specifically to test this hypothesis, suggest that this is precisely what is happening, lending credence to the hypothesis that the shift in strategies across perceptual levels is the element which interacts with the emotional face interference in the FD-e condition.

It could be argued that we could not find an interaction between the M and UD conditions because they, in fact, involve the same processes, as there is some degree of working memory involved even in the M condition, despite the cue stimulus being on screen at all times (i.e., participants may briefly hold the cue face in mind while saccading between and evaluating each the two probe faces to make their match judgment). If that were the case, and similarly there was no difference between the UD and FD-e conditions, then there should be no difference in the encoding of the stimuli between memory conditions. Therefore, there should be no effect of which condition the images used in the long-term recognition memory were drawn from. Irrespective of whether the first encountered memory condition was M, UD, or FD-e, we would expect to see comparable accuracy on the long-term recognition memory task. Our data, on the other hand, show distinct signatures for each process (Fig. 7). Specifically, the higher d’ in the long-term recognition task for stimuli originating from the UD condition indicates a greater recruitment of memory processes and enhanced encoding in this condition relative to the M condition, arguing against the explanation that the lack of interaction stems from an overlap in the processes involved in both UD and M. The lower d’ in the FD-e condition on the other hand, suggests that interference in the face domain also degrades encoding abilities. This decrease in long-term memory abilities might be an additional reflection of the interference in holistic processing in the FD-e condition, as old/new face recognition, which involves comparing very different faces (as opposed to the very similar faces in the match-to-sample parts of the FMP task), is likely to rely predominantly on holistic, rather than feature-based processing.

Unlike all other comparisons across conditions in this study which were all always within the same participants, the comparison of d’ across M, UD, and FD-e involves different participants for each condition (as the initial memory block from which the old images are drawn was randomised across participants). However, as the same result replicated independently across all datasets, and was strongest in the concatenated dataset of nearly 600 participants, this appears to be a robust result. Unfortunately, we did not have enough data from the dataset including the math condition (which had 150 participants and four different memory conditions, so that for each memory condition there were only ∼35 participants for which it appeared first) to reliably calculate how the d’ from the FD-m condition compared to the others. Our hypothesis, however, given the other results, would be that it behaves the same as the UD condition, with better encoding and therefore higher performance on the long-term memory for images drawn from this condition.

Apart from testing differences in the encoding across the three conditions as inferred from the subsequent long-term memory performance, this design allows us to connect our findings to naturalistic face memory in three important ways. First, it allows us to extend our findings from working memory into long-term memory, as evidenced by the moderate correlations we find between the two (Fig. 7e). Second, participants are unaware that they will be asked to perform this task when they start the experiment, and thus aren’t motivated to pay extra attention to the first set of faces. Third, this design reflects real-life scenarios, in which participants encounter many faces which they maintain briefly in memory, out of which only a fraction is essential on longer timescales.

Although our results are consistent and robust, there are aspects of our experimental approach that impose some limitations on the generalisability of the findings. Our face stimulus set is large and diverse and controlled for many low-level visual features, but the to-be-remembered faces were all greyscale. While we believe these results would replicate in more ecologically valid settings, it is possible that different task conditions, types of interference, memory demands, and context might uncover additional interaction patterns. For instance, it is an open question as to whether an active judgement on the emotional faces is necessary to create the observed interaction in the FD-e condition, or whether merely viewing emotional faces in absence of a judgement would be enough to impair performance. This question is difficult to test in online experimental paradigms without eye tracking to confirm that participants attend to the face distractor during the delay period without an overt response. However, given the lack of similar effects in the FD-m and FD-v conditions, which both involve active tasks, it is more likely that these effects result from the nature of the distractor stimuli (faces), rather than the nature of the task. We should also note that we explicitly chose to focus on morphing manipulations for studying perception, rather than different views or images of the same face (as done, for instance, in the Glasgow Face Matching Task). While other types of manipulation would allow us to study higher-order identity representations, it is difficult to disentangle the perceptual processes from other inherent memory processes involved in these tasks, such as those that might be required to match identity across different representations of the same face.

Though it did not affect the core findings presented here, our data revealed other interesting features of face processing, such as the lack of symmetry in the morphing, which should be explored more in depth in future studies. Also, we did not focus in this study on individual differences in performance, although these are clearly visible in our data and have been described before^100,101^. Future neuroimaging studies can harness these individual differences in behaviour between and across the different memory conditions and perceptual levels to search for the neural variance which explains them. Combining fMRI and behavioural testing could help uncover the distinct neural networks involved in the different processes, as well as those shared across conditions.

Our results reinforce the fundamental separation of face processing and face memory, which was our most consistent and replicable finding across all five datasets and 907 participants. That these processes are separable at the most basic level, however, by no means negates the possibility of interactions under various different conditions. Our study explores only a small fraction of the potential cognitive space for such interactions, but our results already highlight certain conditions that are sufficient for interactions and others that are not, and suggest a possible mechanism, which does not necessarily involve overlapping cognitive processes. Though we focus on the face domain, it is not unlikely that these results extend to the broader context of the independence and interactions of perception and memory. Understanding the relationship between these basic processes will help us to better localize functions in the brain, and has far-reaching implications for various memory disorders as well as different agnosias.

## Methods

### Stimuli selection and preparation for FMP task

Face stimuli for all tasks and datasets in this manuscript (excluding the emotional distractor faces in Filled Delay – emotions, FD-e) were selected from several databases to account for maximal diversity in race and age and to avoid any possible artefacts coming from the database selection (for example, previous exposure, quality of pictures, race effect)^102^ and only emotionally neutral faces were used: 3DSK^103^, Bogazici Face Database (BFD)^104^, Karolinska Directed Emotional Faces (KDEF)^105,106^, Chicago Face Database CFD (original^107^, Multi-racial extension^108^, India^109^), Face Research Lab London Set (London)^58^, Oslo^110^, Psychological Image Collection at Stirling (PICS) (http://pics.psych.stir.ac.uk/), RADIATE^111^. The total representation of the databases is shown in SI Table 12. For the experiment, there were always exactly 50% female and 50% male faces and the racial representation was matched for each form to represent the percentage in the full sample of 1640 face stimuli (6 forms of 240 stimuli each), see SI Table 13. For the emotional distractor task on FD-e trials, emotional stimuli from FACES^59^ were used.

All images for stimuli (except the emotional faces) were processed in the following way. First, all face images were cropped, resized and converted to black and white scale in Photoshop CC (Adobe System Inc., CA, USA). All significant birthmarks were removed. Second, all stimuli were standardised to have the same luminance and brightness and an oval mask was added over them using the cv2^112^ package in Python. Then, two images, one for females, and one for males, were selected as templates for morphing. That ensures that all stimuli are morphed towards the same face (concerning gender) and should have the same distance from it, which is comparable across stimuli. Subsequently, 99 landmarks were manually placed on each face outlining the edges of the face, the eyebrows, the eyes, the iris, the mouth, and the nose, and the stimuli were then morphed to be 0%, 24%, 30%, and 36% of the template image using the Abrosoft FantaMorph software (http://www.fantamorph.com/index.html). This software has been widely used in previous studies to create a face series between two expressions^113–115^, dissect face memory and perception development^32^, or evaluate race perception^116^. Because morphing works on the principle of Delaunay triangulation followed by weighted pixel averaging in the corresponding triangles (projective/affine transformation), the procedure yields blurred and fainted-colour images. To remedy this artefact, we sharpened twice all morphed images using the cv2 package in Python. As sharpening also slightly changes the appearance of the face, we sharpened once the original, 0% morph (unmorphed) images as well to make them visually similar to the morphed images. The full procedure is depicted in Supplementary Figure 12. Unlike the face stimuli used for memory conditions, images of emotional distractors were left unmodified, i.e., in colour, and no corrections for brightness or contrast were made.

In Dataset 2, the stimuli were matched in image entropy as much as possible across all nine conditions (memory and perception difficulty). Image entropy is defined as Shannon entropy of an image and was calculated for each image separately using the scikit image package^117^ in Python (https://scikit-image.org/docs/stable/api/skimage.measure.html#skimage.measure.shannon_entropy).

### Validation and selection of food stimuli for Visual interference condition

This was a pilot experiment to select and assess the difficulty of food distractors for the vegetarian classification task in the FD-v condition described above. In this pilot, 90 images of food plates, 45 vegetarian and 45 non-vegetarian, were presented to participants for 4s (like in the Filled Delay condition of the FMP task) after which participants were asked whether the dish was vegetarian or not (Yes/No). In total, N=51 participants (mean age 35.3 ± 11.0, 18 females, 32 males, 1 not stated) were recruited and one was rejected due to technical problems during the task. Participants took on average 15□min to complete the task.

The images were subsequently ordered based on participants’ accuracy (agreement), and we selected the top 30 non-vegetarian and the top 30 vegetarian dishes as stimuli for the full experiment. These stimuli were correctly identified at least by 73% of participants (average classification accuracy (vegetarian/non-vegetarian) 87.5% ± 7% ranging from 73% to 100%).

### Experimental design

The design of this experiment was inspired by Hacker and Biederman^57^ and is shown in Figure 1. Similar to that experiment, the FMP contains two dimensions – two variables whose independence is assessed by examining the interaction between them. The first dimension is the perceptual component and contains three perceptual difficulty levels (Fig. 1a). Different difficulty levels are achieved by morphing all stimuli to a single template face (male faces to a male template and female faces to a female template) and using different morph levels to parametrically change the perceptual difficulty. These levels go from 24% morphs (hardest, level 3), 30% morphs (level 2) to 36% morphs (easiest, level 1), see gallery of example stimuli in Supplementary Figure 13. This morphing method, together with a triangular arrangement, has previously been used in perceptual training for prosopagnostics^118,119^. In the testing phase, both the morphed and unmorphed (original) images are presented, one of which matches the target face which was presented initially. In all datasets except for Dataset 3, the cue stimulus is always the unmorphed face. In Dataset 3, the cue stimulus is either the morphed or the unmorphed face, and participants still must choose the exact match from the two probes (morphed and unmorphed). Each morph level was approximately equally represented in the trials with morphed cue stimuli. The proportion of trials with morphed versus unmorphed cue stimuli was 0.3:0.7 in M, 0.27:0.73 in UD, and 0.24:0.76 in FD-e. Each perceptual level contains 20 stimuli (ten females, ten males), and the difficulty is randomised across trials.

Orthogonal to the perceptual conditions, the basic design of the FMP task utilises three memory conditions (memory conditions) that introduce varying demands on working memory processes (Fig. 1b). The order of memory conditions is randomised across participants, and in each memory condition, participants view 60 stimuli, 20 from each of the perceptual difficulty levels, randomised and split into three chunks to avoid fatigue (Fig. 1c). Participants always complete all three chunks of the condition before moving on to the next task. The easiest condition with little to no demand for working memory is a perceptual face Matching (M) task. In the M condition, participants view a face for 3s, then two other face stimuli (the unmorphed face and a morphed face) appear below it on the screen forming a triangular arrangement, and participants are asked to choose which of the two faces is an exact match to the top face. The cue stimulus remains on the screen for the entire trial duration. Participants have 3s to answer.

In the second memory condition, Unfilled Delay (UD), the presentation of the cue stimulus is followed by a blank screen that is displayed for 4s. Then, only the two probe faces appear, and participants have 3.5s to choose the matching face from the two options without the presence of the cue stimulus.

The third and most challenging memory condition, Filled Delay with emotional interference (FD-e), is analogous to the UD condition but includes active face memory interference. Instead of a blank screen, participants see an emotional face for 4s and must remember the emotion it shows. When the distractor disappears, the two probe faces appear (but not the cue stimulus), and participants are asked to choose which of them is an exact match to the cue stimulus displayed earlier. Finally, two emotion labels appear, and participants must choose the label which most accurately describes the emotion of the distractor image. Due to the different preprocessing of the stimuli (see section above), the distractor face was visually very different from the cue and probe stimuli.

To further test the effect of the nature of the interference on the difference between the Unfilled Delay and Filled Delay conditions, we introduced a Filled Delay with math (FD-m) condition that added orthogonal interference to the face memory process using short arithmetic problems (Dataset 4). Solving arithmetic problems has previously been successful in manipulating memory load^120^. The FD-m condition was structurally similar to the FD-e condition, but presented participants with moderately complex equations, concatenating addition or subtraction with multiplication or division (e.g., 2 * 5 + 43 = 94), instead of faces with emotions. As with the emotional faces in the FD-e condition, equations in the FD-m condition were displayed during the 4s delay period. Participants had to evaluate and remember whether the equation was True or False, and as in the FD-e condition, were asked to respond to this additional question after selecting which of the two probe stimuli was identical to the initially presented cue stimulus. The location (left or right) of the True/False labels on the screen was randomised across trials.

Lastly, to examine whether visual interference alone could account for the interaction observed between UD and FD-e conditions, we introduced a Filled Delay with visual non-face distractor (FD-v) condition using complex, non-face visual stimuli (Dataset 5). The FD-v condition was structurally identical to the FD-e and FD-m conditions, but instead of emotional faces or equations, participants were shown images of food plates during the 4s delay period. Each image was accompanied by a binary categorisation task in which participants had to judge and remember whether the dish was vegetarian or not (Yes/No). As in the FD-e condition, participants responded to this question immediately after selecting which of the two probe stimuli matched the initially presented face. As in FD-m, the location (left or right) of the True/False labels on the screen was randomised across trials.

The FMP design consists of three levels of perception fully crossed with three (or four) levels of memory design, with each condition containing 20 trials, yielding a total of 180 trials (240 trials for the dataset including FD-m and FD-v). The number of trials was tested and optimised for reliability and task duration^121^. Images never repeat across trials. Additionally, the FMP contains a final memory test that appears at the end after all conditions (Fig. 1d).

This Long-term recognition memory condition presents participants with 60 stimuli (30 old, 30 new) for 3s each and asks whether they saw these faces before (Yes/No) and how confident they are (1 – not at all confident, 2 – moderately confident, 3 – very confident). The confidence ratings were not used in our analyses. The 30 old stimuli are selected randomly from the first memory condition the participant encountered (i.e., from the beginning of the experiment, with a retention interval of approximately 30 minutes). As the order of the memory conditions is randomised across participants, this first block could belong to any of the three memory conditions.

Before beginning each task in the main experiment, participants were first shown example screenshots and guided through a single trial for that specific task. They then completed practice trials with feedback – four trials for the first task they encountered and two trials for each subsequent task. This structure (walkthrough, practice, task) was repeated for each task. As the overall task format remained consistent, this approach helped ensure understanding while avoiding unnecessary repetition.

Finally, we conducted an additional control experiment specifically testing the inversion effect across perception difficulty levels. This experiment included only the UD condition. Half of the trials had inverted faces and there were 40 trials per difficulty level per inversion (upright, inverted), yielding a total of 240 trials.

### Data collection

This study was not pre-registered. All data were collected online and participants were recruited using the online platform Prolific (www.prolific.co). To minimise variability in device type and display conditions, only participants using desktop or laptop computers were eligible to take part in the studies. Prior to the task, participants were informed that the experiment would not run on phones or tablets. This restriction was enforced through Prolific’s prescreening options and a PHP-based device check that blocked access for mobile or tablet users. We also recorded technical data including screen resolution, browser, and operating system. While some variation in screen size and viewing conditions may still be present, this is an inherent limitation of online experiments.

All demographic data, including age and sex, were provided by Prolific and the demographic representation of our participants is shown in Supplementary Figure 14. We collected five datasets, each with a slight modification of the paradigm (as detailed below) and different participants, but all datasets were geared towards addressing our key research question.

Participants spent between 40 and 50 minutes completing tasks for Datasets□1,L□2, andL□3, while DatasetL□4 (which included the FD-m condition) took an average of 78 minutes and Dataset□5 (with the FD-v condition) took an average of 75 minutes. The inversion experiment required approximately 53 minutes to finish.

#### Dataset 1

The first dataset was collected in two independent phases. In the first phase, data were collected from 83 participants (mean age 26.2 ± 7.8, 41 females, 42 males) and six were excluded (see exclusion criteria below), leaving N=77 who took on average 40 minutes to complete all tasks. This phase did not include the long-term recognition test at the end of the experiment (Dataset 1a).

In the second phase, the long-term recognition test was added, and data were collected from 194 participants (mean age 32.4 ± 7.6, 100 females, 92 males, 2 not stated) out of which twelve participants were excluded, leaving N=182 (Dataset 1b).

#### Dataset 2 – randomised stimuli matched for entropy

In total, 167 participants were collected for this dataset. Fourteen participants were excluded during preprocessing, leaving N=153 (mean age 31.6 ± 10.1, 86 females, 67 males). For 92 participants, the stimuli were completely randomised across conditions, trials, and participants.

#### Dataset 3 – randomising cue stimuli

A total of 271 participants (mean age 25.4 ± 8.1, 105 females, 165 males, 1 not stated) were collected for this dataset. After applying exclusion criteria, nine participants were excluded and 262 participants remained. Although these participants performed several sessions of the experiment on different days, for this study, we used data only from the first day.

#### Dataset 4 – Math interference

This design contained three perceptual difficulty levels and four memory levels – the regular three described in Figure 1 (M, UD, FD-e) and an additional interference level orthogonal to face processing (Filled Delay with Math, Fig. 4a, described above). A total of 179 participants started the study (mean age 31.1 ± 10.9, 65 females, 98 males, 16 not stated). Twenty-nine were excluded during preprocessing, leaving N=150.

#### Dataset 5 – Visual interference

This design contained three perceptual difficulty levels and four memory levels --the regular three described in Figure 1 (M, UD, FD-e) and an additional interference level orthogonal to face processing with visual stimuli matched in size and complexity to faces (Filled Delay with Visual interference, Fig. 5a). The long-term memory task was not included in this experiment. A total of 99 participants started the study, out of which sixteen were excluded during preprocessing due to the same exclusion criteria as Datasets 1-4, leaving N=83 (mean age 32.5 ± 12.5, 38 females, 45 males).

#### Inversion experiment

This experiment contained three perceptual difficulty levels and one memory level (UD). We recruited 71 participants (mean age 33.4 ± 10.9, 34 females, 35 males, 2 not stated).

### Data quality control and exclusion criteria

To control for the quality of the data, which is especially important when conducting data collection online, we set strict criteria for exclusion. We chose to exclude participants based on a combination of their accuracy, reaction time (RT) and individual trial responses. We assessed the following criteria and excluded participants from a dataset if two or more of the following criteria were met:

- Average RT was 2 standard deviations (SD) faster than the group mean.
- Standard deviation of RT was less than 2 SD below the standard deviation of the group.
- Average sequence length of a single response (i.e., repeatedly indicating the same response across consecutive trials) was 2 SD greater than the group mean sequence length.
- Accuracy below chance (50%) in Matching (M) trials as this would indicate that they were not paying attention during the tasks.

If a participants accuracy was greater than 0.5 SD below the mean, they were included, regardless of their RT and individual trial responses. Additionally, participants with more than 30% missed trials (not answered on time) either in all memory conditions or each of the distractor tasks (emotions, equations) were inspected individually and excluded if the pattern of responses suggested they were not paying attention to the task.

### Bayesian modelling

To analyse the relationship between memory conditions, perceptual difficulty, and accuracy, we employed a Bayesian mixed-effects modelling approach. The model fitting was performed using the Bambi^61^ and PyMC^122^ and visualised using arviz^123^ packages in Python. We fitted two hierarchical models, differing in the included interaction term:

Interaction model:

1. y ∼ condition * difficulty + (1 | participant)

interaction model:

2. y ∼ condition + difficulty + (1 | participant)

where *y* is the mean accuracy on 20 trials per participant in a given combination of memory condition and perceptual difficulty; *condition* and *difficulty* are fixed effects corresponding to memory and perceptual difficulties, respectively; and *(1 | participant)* accounts for random intercepts at the participant level. All variables were treated as categorical. The Gaussian family with an identity link was used and default, weakly informative Bambi priors estimated from the data were chosen and checked visually before the model was fitted (for details, see SI).

We omitted data points with missing values and fit our models in parallel using four chains with PyMC’s adaptive Hamiltonian Monte Carlo (HMC) algorithm (No-U-Turn Sampler, NUTS). Each chain drew 1,000 samples from the posterior after a minimum of 1,000 burn-in samples. We checked convergence for all independent variables using the Gelman–Rubin statistic r□, ensuring all values were below 1.01, indicating good convergence. We report the 94% high-density intervals (HDI) of the posterior distribution to quantify uncertainty. We considered a fixed effect or interaction significant if its 94% HDI did not include zero following recent practices^60^ and as is the default in the Bambi package^61^.

For model comparison and validation, we used log-likelihood estimation. To test for interactions, we employed two strategies: 1) exploring posterior distributions of interaction parameters, and 2) comparing alternative and null hypothesis models described above. We compared models by estimating the expected log-wise predictive density (ELPD) and standard error (SE) using Pareto-smoothed importance sampling leave-one-out cross-validation^124^ (PSIS-LOO). A higher ELPD indicates a better out-of-sample predictive fit. PSIS-LOO offers advantages, including robustness to weak priors, invariance to parameterisation, and avoiding issues like sensitivity to priors found in Bayes factors.

### Software and used packages

All the analyses were performed using Python 3^125^ and the following packages:

numpy^126^, pandas^127^, matplotlib^128^, seaborn^129^, geopandas^130^, cv2^112^, scipy^131^. Final figures were created using the CanD^132^ package. The Bayesian model fitting was performed using the Bambi^61^ and PyMC^122^ and visualised using arviz^123^ packages.

All the experiments were coded using lab.js^133^ (www.lab.js.org) and run on our servers.

Images were processed in Photoshop CC (Adobe System Inc., CA, USA) and morphed using the Abrosoft FantaMorph software (http://www.fantamorph.com/index.html).

### Code and data availability

Code and data will be made accessible on GitHub and ZENODO at the time of publication.

## Acknowledgements

We would like to thank all participants that took part in this study. We thank Sasha Devore for insightful comments during the writing process. J.K. would like to thank Dana Yakobi and Ron Rotkopf for their advice on statistics, Jan Soukup for advice on mathematical modelling and interpretation, and Anna Uzonyi for fruitful discussions about the design and results of the experiments and tips throughout the entire process. J.K. would also like to thank Shakked Ganor and is extremely grateful for her help with manually annotating face landmarks. This work was generously supported by the European Research Council (ERC-2022-StG 101077921), Israel Science Foundation grant 829/22, the UCLA/Weizmann Collaboration in Neuroscience grant, and the Zuckerman STEM leadership program. M.R. is the incumbent of the Roel C. Buck Career Development Chair. C.R.W. was funded by the National Science Foundation Graduate Research Fellowship Program under Grant No. DGE-2034835. The funders had no role in study design, data collection and analysis, decision to publish or preparation of the manuscript.

## Author Contributions

Jan Kadlec, Catherine Walsh, Jesse Rissman and Michal Ramot conceptualised the study and the task design. Jan Kadlec and Catherine Walsh collected the empirical data. Jan Kadlec analysed and visualised the data. Jan Kadlec and Michal Ramot conceptualised the article and created its framework. Jan Kadlec and Michal Ramot wrote the first draft. All authors contributed to writing and editing the article. All authors participated in data interpretation. Jesse Rissman and Michal Ramot acquired the necessary funding.

## References

1. Scoville, W. B. & Milner, B. Loss of Recent Memory After Bilateral Hippocampal Lesions. J. Neurol. Neurosurg. Psychiatry 20, 11–21 (1957).

2. Mishkin, M., Ungerleider, L. G. & Macko, K. A. Object vision and spatial vision: two cortical pathways. Trends Neurosci. 6, 414–417 (1983).

3. Squire, L. R. & Zola-Morgan, S. The Medial Temporal Lobe Memory System. Science 253, 1380– 1386 (1991).

4. Cosmides, L. & Tooby, J. Beyond intuition and instinct blindness: toward an evolutionarily rigorous cognitive science. Cognition 50, 41–77 (1994).

5. Kanwisher, N., Chun, M. M., McDermott, J. & Ledden, P. J. Functional imaging of human visual recognition. *Cogn*. Brain Res. 5, 55–67 (1996).

6. Bodamer, J. Die Prosop-Agnosie. Arch. Für Psychiatr. Nervenkrankh. 179, 6–53 (1947).

7. Shelton, P. A., Bowers, D., Duara, R. & Heilman, K. M. Apperceptive Visual Agnosia: A Case Study. Brain Cogn. 25, 1–23 (1994).

8. Vecera, S. P. & Gilds, K. S. What Processing Is Impaired in Apperceptive Agnosia? Evidence from Normal Subjects. J. Cogn. Neurosci. 10, 568–580 (1998).

9. Davies-Thompson, J., Pancaroglu, R. & Barton, J. Acquired prosopagnosia: structural basis and processing impairments. Front. Biosci.-Elite 6, 159–174 (2014).

10. Busigny, T., Joubert, S., Felician, O., Ceccaldi, M. & Rossion, B. Holistic perception of the individual face is specific and necessary: Evidence from an extensive case study of acquired prosopagnosia. Neuropsychologia 48, 4057–4092 (2010).

11. Cowell, R. A., Barense, M. D. & Sadil, P. S. A Roadmap for Understanding Memory: Decomposing Cognitive Processes into Operations and Representations. eNeuro 6, (2019).

12. Martin, C. B. & Barense, M. D. Perception and Memory in the Ventral Visual Stream and Medial Temporal Lobe. Annu. Rev. Vis. Sci. 9, 409–434 (2023).

13. Khan, Z. U., Martín-Montañez, E. & Baxter, M. G. Visual perception and memory systems: from cortex to medial temporal lobe. Cell. Mol. Life Sci. 68, 1737–1754 (2011).

14. Megla, E. & Bainbridge, W. Interaction of Perception and Memory. Preprint at 10.31234/osf.io/m4uyt (2023).

15. Tsao, D. Y. & Livingstone, M. S. Mechanisms of Face Perception. Annu. Rev. Neurosci. 31, 411–437 (2008).

16. Oosterhof, N. N. & Todorov, A. The functional basis of face evaluation. Proc. Natl. Acad. Sci. 105, 11087–11092 (2008).

17. Rule, N. O., Krendl, A. C., Ivcevic, Z. & Ambady, N. Accuracy and consensus in judgments of trustworthiness from faces: Behavioral and neural correlates. J. Pers. Soc. Psychol. 104, 409–426 (2013).

18. Sussman, A. B., Petkova, K. & Todorov, A. Competence ratings in US predict presidential election outcomes in Bulgaria. J. Exp. Soc. Psychol. 49, 771–775 (2013).

19. Oruc, I., Balas, B. & Landy, M. S. Face perception: A brief journey through recent discoveries and current directions. Vision Res. 157, 1–9 (2019).

20. Willis, J. & Todorov, A. First Impressions: Making Up Your Mind After a 100-Ms Exposure to a Face. Psychol. Sci. 17, 592–598 (2006).

21. Arioli, M., Crespi, C. & Canessa, N. Social Cognition through the Lens of Cognitive and Clinical Neuroscience. BioMed Res. Int. 2018, 4283427 (2018).

22. Smith, M. L., Cottrell, G. W., Gosselin, F. & Schyns, P. G. Transmitting and Decoding Facial Expressions. Psychol. Sci. 16, 184–189 (2005).

23. Duchaine, B. & Yovel, G. A Revised Neural Framework for Face Processing. Annu. Rev. Vis. Sci. 1, 393–416 (2015).

24. Ramot, M., Walsh, C. & Martin, A. Multifaceted Integration: Memory for Faces Is Subserved by Widespread Connections between Visual, Memory, Auditory, and Social Networks. J. Neurosci. 39, 4976–4985 (2019).

25. Kanwisher, N., McDermott, J. & Chun, M. M. The Fusiform Face Area: A Module in Human Extrastriate Cortex Specialized for Face Perception. J. Neurosci. 17, 4302–4311 (1997).

26. Haxby, J. V., Hoffman, E. A. & Gobbini, I. M. The distributed human neural system for face perception. Trends Cogn. Sci. 4, 223–233 (2000).

27. Gobbini, M. I. & Haxby, J. V. Neural systems for recognition of familiar faces. Neuropsychologia 45, 32–41 (2007).

28. Kriegeskorte, N., Formisano, E., Sorger, B. & Goebel, R. Individual faces elicit distinct response patterns in human anterior temporal cortex. Proc. Natl. Acad. Sci. 104, 20600–20605 (2007).

29. Rajimehr, R., Young, J. C. & Tootell, R. B. H. An anterior temporal face patch in human cortex, predicted by macaque maps. Proc. Natl. Acad. Sci. 106, 1995–2000 (2009).

30. Kreiman, G., Koch, C. & Fried, I. Category-specific visual responses of single neurons in the human medial temporal lobe. Nat. Neurosci. 3, 946–953 (2000).

31. Chen, Y. Y., Areti, A., Yoshor, D. & Foster, B. L. Perception and Memory Reinstatement Engage Overlapping Face-Selective Regions within Human Ventral Temporal Cortex. J. Neurosci. 44, (2024).

32. Weigelt, S. et al. Domain-specific development of face memory but not face perception. Dev. Sci. 17, 47–58 (2014).

33. Ofen, N. & Shing, Y. L. From perception to memory: Changes in memory systems across the lifespan. Neurosci. Biobehav. Rev. 37, 2258–2267 (2013).

34. Stantić, M., Hearne, B., Catmur, C. & Bird, G. Use of the Oxford face matching test reveals an effect of ageing on face perception but not face memory. Cortex 145, 226–235 (2021).

35. Barton, J. J. S., Davies-Thompson, J. & Corrow, S. L. Chapter 10 -Prosopagnosia and disorders of face processing. in Handbook of Clinical Neurology (eds. Barton, J. J. S. & Leff, A.) vol. 178 175–193 (Elsevier, 2021).

36. Cook, R. & Biotti, F. Developmental prosopagnosia. Curr. Biol. 26, R312–R313 (2016).

37. Corrow, S. L., Dalrymple, K. A. & Barton, J. J. Prosopagnosia: current perspectives. Eye Brain 8, 165–175 (2016).

38. Bate, S. & Tree, J. J. The Definition and Diagnosis of Developmental Prosopagnosia. Q. J. Exp. Psychol. 70, 193–200 (2017).

39. Duchaine, B. & Nakayama, K. Dissociations of Face and Object Recognition in Developmental Prosopagnosia. J. Cogn. Neurosci. 17, 249–261 (2005).

40. Duchaine, B. C. & Nakayama, K. Developmental prosopagnosia: a window to content-specific face processing. Curr. Opin. Neurobiol. 16, 166–173 (2006).

41. Rossion, B. Twenty years of investigation with the case of prosopagnosia PS to understand human face identity recognition. Part I: Function. Neuropsychologia 108278 (2022) doi:10.1016/j.neuropsychologia.2022.108278.

42. Rossion, B. Understanding face perception by means of prosopagnosia and neuroimaging. Front. Biosci.-Elite 6, 258–307 (2014).

43. De Renzi, E., Faglioni, P., Grossi, D. & Nichelli, P. Apperceptive and associative forms of prosopagnosia. Cortex J. Devoted Study Nerv. Syst. Behav. 27, 213–221 (1991).

44. Troup, L. J., Bastidas, S., Nomi, J. S., Nguyen, M. T. & Tong, T. Face and emotional expression processing and event-related potentials in a case study of impaired face perception. Edorium J. Psychol. 1, 9–17 (2015).

45. Tippett, L. J., Miller, L. A. & Farah, M. J. Prosopamnesia: a selective impairment in face learning. Cognitive Neuropsychology 17, 241–255 (2000).

46. Barton, J. J. Disorders of face perception and recognition. Neurol. Clin. 21, 521–548 (2003).

47. Smith, C. N. et al. When recognition memory is independent of hippocampal function. Proc. Natl. Acad. Sci. 111, 9935–9940 (2014).

48. Wingrove, J. R. B. & Tree, J. J. Can face recognition be selectively preserved in some cases of amnesia? A cautionary tale. Cortex 173, 283–295 (2024).

49. Shrager, Y., Gold, J. J., Hopkins, R. O. & Squire, L. R. Intact Visual Perception in Memory-Impaired Patients with Medial Temporal Lobe Lesions. J. Neurosci. 26, 2235–2240 (2006).

50. Rossion, B. Damasio’s error – Prosopagnosia with intact within-category object recognition. J. Neuropsychol. 12, 357–388 (2018).

51. Rossion, B. Prosopdysgnosia? What could it tell us about the neural organization of face and object recognition? Cogn. Neuropsychol. 35, 98–101 (2018).

52. Haxby, J. V. et al. Face encoding and recognition in the human brain. Proc. Natl. Acad. Sci. 93, 922–927 (1996).

53. Liu, X., Li, X., Song, Y. & Liu, J. Separate and Shared Neural Basis of Face Memory and Face Perception in Developmental Prosopagnosia. Front. Behav. Neurosci. 15, 668174 (2021).

54. Haeger, A. et al. Face Processing in Developmental Prosopagnosia: Altered Neural Representations in the Fusiform Face Area. Front. Behav. Neurosci. 15, (2021).

55. Manippa, V., Palmisano, A., Ventura, M. & Rivolta, D. The Neural Correlates of Developmental Prosopagnosia: Twenty-Five Years on. Brain Sci. 13, 1399 (2023).

56. Stantic, M. et al. The Oxford Face Matching Test: A non-biased test of the full range of individual differences in face perception. Behav. Res. Methods 54, 158–173 (2022).

57. Hacker, C. & Biederman, I. The Proficiency for Distinguishing Faces Is Independent of the Proficiency for Remembering Them. https://osf.io/9bwct (2019) doi:10.31234/osf.io/9bwct.

58. DeBruine, L. & Jones, B. Face Research Lab London Set. Figshare (2017) doi:10.6084/m9.figshare.5047666.v3.

59. Ebner, N. C., Riediger, M. & Lindenberger, U. FACES—A database of facial expressions in young, middle-aged, and older women and men: Development and validation. Behav. Res. Methods 42, 351–362 (2010).

60. Gehmacher, Q. et al. Eye movements track prioritized auditory features in selective attention to natural speech. Nat. Commun. 15, 3692 (2024).

61. Capretto, T. et al. Bambi: A Simple Interface for Fitting Bayesian Linear Models in Python. J. Stat. Softw. 103, 1–29 (2022).

62. Sweller, J. Cognitive Load During Problem Solving: Effects on Learning. Cogn. Sci. 12, 257– 285 (1988).

63. Sweller, J. CHAPTER TWO -Cognitive Load Theory. in Psychology of Learning and Motivation (eds. Mestre, J. P. & Ross, B. H.) vol. 55 37–76 (Academic Press, 2011).

64. Schneider, W. & Shiffrin, R. M. Controlled and automatic human information processing: I. Detection, search, and attention. Psychol. Rev. 84, 1–66 (1977).

65. Richler, J. J., Tanaka, J. W., Brown, D. D. & Gauthier, I. Why does selective attention to parts fail in face processing? J. Exp. Psychol. Learn. Mem. Cogn. 34, 1356–1368 (2008).

66. Valentine, T. Upside-down faces: A review of the effect of inversion upon face recognition. Br. J. Psychol. 79, 471–491 (1988).

67. Richler, J. J., Mack, M. L., Palmeri, T. J. & Gauthier, I. Inverted faces are (eventually) processed holistically. Vision Res. 51, 333–342 (2011).

68. Rossion, B. Picture-plane inversion leads to qualitative changes of face perception. Acta Psychol. (Amst.) 128, 274–289 (2008).

69. Piepers, D. & Robbins, R. A Review and Clarification of the Terms “holistic,” “configural,” and “relational” in the Face Perception Literature. Front. Psychol. 3, (2012).

70. Klargaard, S. K., Starrfelt, R., Petersen, A. & Gerlach, C. Topographic processing in developmental prosopagnosia: Preserved perception but impaired memory of scenes. Cogn. Neuropsychol. 33, 405–413 (2016).

71. Chai, W. J., Abd Hamid, A. I. & Abdullah, J. M. Working Memory From the Psychological and Neurosciences Perspectives: A Review. Front. Psychol. 9, (2018).

72. Christophel, T. B., Klink, P. C., Spitzer, B., Roelfsema, P. R. & Haynes, J.-D. The Distributed Nature of Working Memory. Trends Cogn. Sci. 21, 111–124 (2017).

73. Rissman, J. & Wagner, A. D. Distributed Representations in Memory: Insights from Functional Brain Imaging. Annu. Rev. Psychol. 63, 101–128 (2012).

74. Cohen, J. D. et al. Temporal dynamics of brain activation during a working memory task. Nature 386, 604–608 (1997).

75. Funahashi, S. Working Memory in the Prefrontal Cortex. Brain Sci. 7, 49 (2017).

76. Gazzaley, A., Rissman, J. & D’esposito, M. Functional connectivity during working memory maintenance. Cogn. Affect. Behav. Neurosci. 4, 580–599 (2004).

77. Rissman, J., Gazzaley, A. & D’Esposito, M. Dynamic adjustments in frontal, hippocampal, and inferior temporal interactions with increasing visual working memory load. Cereb. Cortex N. Y. N 1991 18, 1618–1629 (2008).

78. Wilf, M. et al. Using virtual reality-based neurocognitive testing and eye tracking to study naturalistic cognitive-motor performance. Neuropsychologia 194, 108744 (2024).

79. Al-Yahya, E. et al. Cognitive motor interference while walking: A systematic review and meta-analysis. Neurosci. Biobehav. Rev. 35, 715–728 (2011).

80. Baddeley, A. D. & Hitch, G. Working Memory. in Psychology of Learning and Motivation (ed. Bower, G. H.) vol. 8 47–89 (Academic Press, 1974).

81. Biehl, S. C. et al. The impact of task relevance and degree of distraction on stimulus processing. BMC Neurosci. 14, 107 (2013).

82. Sreenivasan, K. K. & Jha, A. P. Selective Attention Supports Working Memory Maintenance by Modulating Perceptual Processing of Distractors. J. Cogn. Neurosci. 19, 32–41 (2007).

83. Costa, M. et al. Aversive memory formation in humans involves an amygdala-hippocampus phase code. Nat. Commun. 13, 6403 (2022).

84. Sergerie, K., Chochol, C. & Armony, J. L. The role of the amygdala in emotional processing: A quantitative meta-analysis of functional neuroimaging studies. Neurosci. Biobehav. Rev. 32, 811– 830 (2008).

85. van den Bulk, B. G. et al. Amygdala activation during emotional face processing in adolescents with affective disorders: the role of underlying depression and anxiety symptoms. Front. Hum. Neurosci. 8, 393 (2014).

86. Bradley, M. M. & Sambuco, N. Emotional Memory and Amygdala Activation. Front. Behav. Neurosci. 16, (2022).

87. Paré, D. & Headley, D. B. The amygdala mediates the facilitating influence of emotions on memory through multiple interacting mechanisms. Neurobiol. Stress 24, 100529 (2023).

88. Farah, M. J., Wilson, K. D., Drain, M. & Tanaka, J. N. What is ‘special’ about face perception? Psychol. Rev. 105, 482–498 (1998).

89. Leong, B. Q. Z., Estudillo, A. J. & Hussain Ismail, A. M. Holistic and featural processing’s link to face recognition varies by individual and task. Sci. Rep. 13, 16869 (2023).

90. Boutet, I., Gentes-Hawn, A. & Chaudhuri, A. The influence of attention on holistic face encoding. Cognition 84, 321–341 (2002).

91. Yin, R. K. Looking at upside-down faces. J. Exp. Psychol. 81, 141–145 (1969).

92. McKone, E. & Yovel, G. Why does picture-plane inversion sometimes dissociate perception of features and spacing in faces, and sometimes not? Toward a new theory of holistic processing. Psychon. Bull. Rev. 16, 778–797 (2009).

93. Liu, X. & Tanaka, J. W. Holistic perception of faces in 17 milliseconds: Evidence from three measures. J. Vis. 19, 92 (2019).

94. Tanaka, J. W. & Farah, M. J. Parts and Wholes in Face Recognition. Q. J. Exp. Psychol. Sect. A 46, 225–245 (1993).

95. Rakover, S. S. Featural vs. configurational information in faces: A conceptual and empirical analysis. Br. J. Psychol. 93, 1–30 (2002).

96 . Richler, J. J., Cheung, O. S. & Gauthier, I. Holistic Processing Predicts Face Recognition. Psychol. Sci. 22, 464–471 (2011).

97. Lee, J. K. W., Janssen, S. M. J. & Estudillo, A. J. A featural account for own-face processing? Looking for support from face inversion, composite face, and part-whole tasks. -Percept. 13, 20416695221111409 (2022).

98. Rezlescu, C., Susilo, T., Wilmer, J. B. & Caramazza, A. The inversion, part-whole, and composite effects reflect distinct perceptual mechanisms with varied relationships to face recognition. J. Exp. Psychol. Hum. Percept. Perform. 43, 1961–1973 (2017).

99. Maurer, D., Grand, R. L. & Mondloch, C. J. The many faces of configural processing. Trends Cogn. Sci. 6, 255–260 (2002).

100. Fysh, M. C. Individual differences in the detection, matching and memory of faces. Cogn. Res. Princ. Implic. 3, 20 (2018).

101. Wilmer, J. B. Individual Differences in Face Recognition: A Decade of Discovery. Curr. Dir. Psychol. Sci. 26, 225–230 (2017).

102. Valentine, T. A Unified Account of the Effects of Distinctiveness, Inversion, and Race in Face Recognition. Q. J. Exp. Psychol. Sect. A 43, 161–204 (1991).

103. DeBruine, L. & Jones, B. C. 3DSK face set with webmorph templates. Open Sci. Framew. (2020) doi:10.17605/OSF.IO/A3947.

104. Saribay, S. A. et al. The Bogazici face database: Standardized photographs of Turkish faces with supporting materials. PLOS ONE 13, e0192018 (2018).

105. Lundqvist, D., Flykt, A., & Öhman, A. The Karolinska Directed Emotional Faces—KDEF. Karolinska Inst. (1998) doi:10.1037/t27732-000.

106. Garrido, M. V. & Prada, M. KDEF-PT: Valence, Emotional Intensity, Familiarity and Attractiveness Ratings of Angry, Neutral, and Happy Faces. Front. Psychol. 0, (2017).

107. Ma, D. S., Correll, J. & Wittenbrink, B. The Chicago face database: A free stimulus set of faces and norming data. Behav. Res. Methods 47, 1122–1135 (2015).

108. Ma, D. S., Kantner, J. & Wittenbrink, B. Chicago Face Database: Multiracial expansion. Behav. Res. Methods (2020) doi:10.3758/s13428-020-01482-5.

109. Lakshmi, A., Wittenbrink, B., Correll, J. & Ma, D. S. The India Face Set: International and Cultural Boundaries Impact Face Impressions and Perceptions of Category Membership. Front. Psychol. 12, (2021).

110. Chelnokova, O. et al. Rewards of beauty: the opioid system mediates social motivation in humans. Mol. Psychiatry 19, 746–747 (2014).

111. Conley, M. I. et al. The racially diverse affective expression (RADIATE) face stimulus set. Psychiatry Res. 270, 1059–1067 (2018).

112. OpenCV. Open Source Computer Vision Library. (2015).

113. Fox, C. J. & Barton, J. J. S. What is adapted in face adaptation? The neural representations of expression in the human visual system. Brain Res. 1127, 80–89 (2007).

114. Butler, A., Oruc, I., Fox, C. J. & Barton, J. J. S. Factors contributing to the adaptation aftereffects of facial expression. Brain Res. 1191, 116–126 (2008).

115. Qiu, S. & Mei, G. Spontaneous recovery of adaptation aftereffects of natural facial categories. Vision Res. 188, 202–210 (2021).

116. Ma, D. S., Kantner, J., Benitez, J. & Dunn, S. Are Morphs a Valid Substitute for Real Multiracial Faces in Race Categorization Research? Pers. Soc. Psychol. Bull. 0146167221989836 (2021) doi:10.1177/0146167221989836.

117. Walt, S. van der et al. scikit-image: image processing in Python. PeerJ 2, e453 (2014).

118. Bate, S. et al. Rehabilitation of face-processing skills in an adolescent with prosopagnosia: Evaluation of an online perceptual training programme. Neuropsychol. Rehabil. 25, 733–762 (2015).

119. Corrow, S. L. et al. Training face perception in developmental prosopagnosia through perceptual learning. Neuropsychologia 134, 107196 (2019).

120. Van Dillen, L. F. & Koole, S. L. How automatic is “automatic vigilance”? The role of working memory in attentional interference of negative information. Cogn. Emot. 23, 1106–1117 (2009).

121. Kadlec, J. A measure of reliability convergence to select and optimize cognitive tasks for individual differences research -Code at the time of final submission. Zenodo 10.5281/zenodo.11564064 (2024).

122. Abril-Pla, O. et al. PyMC: a modern, and comprehensive probabilistic programming framework in Python. PeerJ Comput. Sci. 9, e1516 (2023).

123. Kumar, R., Carroll, C., Hartikainen, A. & Martín, O. A. ArviZ a unified library for exploratory analysis of Bayesian models in Python. (2019) doi:10.21105/joss.01143.

124. Vehtari, A., Gelman, A. & Gabry, J. Practical Bayesian model evaluation using leave-one-out cross-validation and WAIC. Stat. Comput. 27, 1413–1432 (2017).

125. Van Rossum, G. & Drake, F. L. Python 3 Reference Manual. (CreateSpace, Scotts Valley, CA, 2009).

126. Harris, C. R. et al. Array programming with NumPy. Nature 585, 357–362 (2020).

127. McKinney, W. Data Structures for Statistical Computing in Python. in 56–61 (Austin, Texas, 2010). doi:10.25080/Majora-92bf1922-00a.

128. Hunter, J. D. Matplotlib: A 2D Graphics Environment. Comput. Sci. Eng. 9, 90–95 (2007).

129. Waskom, M. L. seaborn: statistical data visualization. J. Open Source Softw. 6, 3021 (2021).

130. Jordahl, K. et al. geopandas/geopandas: v0.10.2. Zenodo 10.5281/zenodo.5573592 (2021).

131. Virtanen, P. et al. SciPy 1.0: fundamental algorithms for scientific computing in Python. Nat. Methods 17, 261–272 (2020).

132. Shinn, M. CanD features. (2022).

133. Henninger, F., Shevchenko, Y., Mertens, U. K., Kieslich, P. J. & Hilbig, B. E. lab.js: A free, open, online study builder. Behav. Res. Methods 54, 556–573 (2022).

134. Psychological Image Collection at Stirling (PICS). http://pics.psych.stir.ac.uk/ (2008).

