## Supplementary Information for "Behavioural separation of face memory and face perception"

##### SI Methods

##### Selecting perceptual difficulty levels

To decide which morph levels to use, we piloted two blocks of the Matching (M) condition (minimal to no memory condition; see Fig. 1a for details) with perceptual levels 14, 20, and 26 percent of the template image. We recruited 59 participants (mean age 42.3 ± 12.9, 22 females, 37 males) who took on average 18 min to complete the task. The average accuracy for difficulty levels of 14, 20, and 26 was 0.625, 0.701, and 0.799, respectively. Since this was supposed to be the easiest condition on the memory scale, we decided to make the perceptual levels even easier to avoid floor effects. We aimed for an average accuracy on the easiest memory and perceptual level of ∼90%. That resulted in choosing levels 24, 30, and 36 as previously described.

##### Priors selection

The Gaussian family with an identity link was used and default, weakly informative Bambi priors estimated from the data were chosen and checked visually before the model was fitted. The following priors were used:

Common-level effects:

- Intercept ~ Normal(mu: 0.7674, sigma: 0.7465)
- condition ~ Normal(mu: [0, 0], sigma: [0.8463, 0.8474])
- difficulty ~ Normal(mu: [0, 0], sigma: [0.8467, 0.8467])
- condition:difficulty ~ Normal(mu: [0, 0, 0, 0], sigma: [1.2691, 1.2718])

Group-level effects:

- 1 | participant ~ Normal(mu: 0.0, sigma: HalfNormal(sigma: 0.7465))

Auxiliary parameters

- Sigma ~ HalfStudentT(nu: 4.0, sigma: 0.1596)

To estimate the effect of cue stimulus, we fitted the following equation:

1. y ~ condition * difficulty * cue_stimulus + (1 | participant)

The same definition was used also in this case with the following priors:

Common-level effects

- Intercept ~ Beta(alpha: 5.0, beta: 2.0)
- condition ~ Normal(mu: 0.0, sigma: 0.1)
- difficulty ~ Normal(mu: 0.0, sigma: 0.1)
- condition:difficulty ~ Normal(mu: 0.0, sigma: 0.1)
- cue_stimulus ~ Normal(mu: 0.0, sigma: 1.0503)
- condition:cue_stimulus ~ Normal(mu: [0., 0.], sigma: [1.4055, 1.4163])
- difficulty:cue_stimulus ~ Normal(mu: [0., 0.], sigma: [1.4091, 1.4091])
- condition:difficulty:cue_stimulus ~ Normal(mu: [0., 0., 0., 0.], sigma: [2.2856, 2.2856, 2.3064, 2.3064])

Group-level effects

- 1 | userID ~ Normal(mu: 0.0, sigma: HalfNormal(sigma: 0.9147))

Auxiliary parameters

- sigma ~ HalfStudentT(nu: 4.0, sigma: 0.2101)

### SI Tables

#### Significance testing using ANOVAs

##### Dataset 1

| Within group factor | Degrees of freedom (numerator) | Degrees of freedom (denominator) | F-value | Uncorrected p-value | Generalised eta-square effect size |
| --- | --- | --- | --- | --- | --- |
| condition | 2 | 512 | 398.61 | 4.1$\cdot$10^-105^ | 0.28 |
| difficulty | 2 | 512 | 159.77 | 1.2$\cdot$10^-54^ | 0.06 |
| condition*difficulty | 4 | 1024 | 21.46 | 5.2$\cdot$10^-17^ | 0.02 |

**Supplementary Table 1.** Summary table of two-way repeated measures ANOVA showing significant main effect of memory and perceptual difficulty and significant interaction of these two effects in Dataset 1.

##### Dataset 2: Randomised stimuli matched for entropy

| Within group factor | Degrees of freedom (numerator) | Degrees of freedom (denominator) | F-value | Uncorrected p-value | Generalised eta-square effect size |
| --- | --- | --- | --- | --- | --- |
| condition | 2 | 304 | 155.65 | 2.9$\cdot$10^-47^ | 0.20 |
| difficulty | 2 | 304 | 94.71 | 1.1$\cdot$10^-32^ | 0.06 |
| condition*difficulty | 4 | 608 | 4.96 | 6.1$\cdot$10^-4^ | 0.006 |

**Supplementary Table 2.** Summary table of two-way repeated measures ANOVA showing significant main effect of memory and perceptual difficulty and significant interaction of these two effects in Dataset 2.

##### Dataset 3: Randomising cue stimulus

| Within group factor | Degrees of freedom (numerator) | Degrees of freedom (denominator) | F-value | Uncorrected p-value | Generalised eta-square effect size |
| --- | --- | --- | --- | --- | --- |
| condition | 2 | 512 | 358.24 | 5.0$\cdot$10^-98^ | 0.26 |
| difficulty | 2 | 512 | 224.50 | 9.9$\cdot$10^-71^ | 0.10 |
| condition*difficulty | 4 | 1024 | 14.02 | 3.8$\cdot$10^-11^ | 0.01 |

**Supplementary Table 3.** Summary table of two-way repeated measures ANOVA showing significant main effect of memory and perceptual difficulty and significant interaction of these two effects in Dataset 3.

##### Dataset 4: Math interference

| Within group factor | Degrees of freedom (numerator) | Degrees of freedom (denominator) | F-value | Uncorrected p-value | Generalised eta-square effect size |
| --- | --- | --- | --- | --- | --- |
| condition | 3 | 447 | 109.59 | 3.4$\cdot$10^-53^ | 0.17 |
| difficulty | 2 | 298 | 120.45 | 4.6$\cdot$10^-39^ | 0.06 |
| condition*difficulty | 6 | 894 | 1.48 | 0.18 | 0.002 |

**Supplementary Table 4.** Summary table of two-way repeated measures ANOVA showing significant main effect of memory and perceptual difficulty but no significant interaction of these two effects in Dataset 4.

##### Dataset 5: Visual interference

| Within group factor | Degrees of freedom (numerator) | Degrees of freedom (denominator) | F-value | Uncorrected p-value | Generalised eta-square effect size |
| --- | --- | --- | --- | --- | --- |
| condition | 3 | 246 | 52.31 | 3.5$\cdot$10^-26^ | 0.12 |
| difficulty | 2 | 164 | 58.00 | 8.9$\cdot$10^-20^ | 0.06 |
| condition*difficulty | 6 | 492 | 2.14 | 0.047 | 0.06 |

**Supplementary Table 5.** Summary table of two-way repeated measures ANOVA showing significant main effect of memory and perceptual difficulty and significant interaction of these two effects in Dataset 5.

##### Concatenated datasets 1, 2, and 3

| Within group factor | Degrees of freedom (numerator) | Degrees of freedom (denominator) | F-value | Uncorrected p-value | Generalised eta-square effect size |
| --- | --- | --- | --- | --- | --- |
| condition | 2 | 1332 | 894.83 | 4.6$\cdot$10^-247^ | 0.25 |
| difficulty | 2 | 1332 | 474.11 | 1.9$\cdot$10^-164^ | 0.07 |
| condition*difficulty | 4 | 2664 | 33.01 | 6.6$\cdot$10^-27^ | 0.01 |

**Supplementary Table 6.** Summary table of two-way repeated measures ANOVA showing significant main effect of memory and perceptual difficulty and significant interaction of these two effects in concatenated datasets 1, 2, and 3.

##### Learning / Exposure

We performed three one-way ANOVAs, one per each memory condition, with the order in which this condition appeared as the between-subject factor and accuracy as the dependent variable. The results showed a significant effect of the first condition for the M condition (F(2, 671) = 3.20, p = 0.041; $\eta_{p}^{2}$ = 0.0094) but no difference for UD and FD-e (UD: F(2, 671) = 2.19, p = 0.11; $\eta_{p}^{2}$ = 0.0065; FD-e: F(2, 664) = 2.12, p = 0.12; $\eta_{p}^{2}$ = 0.0063). Subsequent pairwise Tukey-HSD post-hoc tests for the M condition, summarised in SI Table 7 below, show a significant difference between the condition appearing first and second but not first and last or second and last.

| Group 1 | Group 2 | Difference in means | Standard error | T-value | Tukey-HSD corrected p-values | Cohen’s d |
| --- | --- | --- | --- | --- | --- | --- |
| first | last | 0.014 | 0.007 | 1.96 | 0.12 | 0.19 |
| first | second | 0.017 | 0.007 | 2.35 | 0.049 | 0.23 |
| last | second | 0.003 | 0.007 | 0.43 | 0.90 | 0.04 |

**Supplementary Table 7.** Summary table of post-hoc testing of learning in the M condition in concatenated datasets 1, 2, 3.

##### Dataset 1b – d’

We performed one-way ANOVA with the first condition as the between-subject factor and d’ as the dependent variable that showed a significant effect of the first condition (F(2, 179) = 7.72; p = 0.0006; MSE = 1.57; $\eta_{p}^{2}$= 0.079). Subsequently, we performed a pairwise Tukey-HSD post-hoc test and we summarise the results in SI Table 8 below.

| Group 1 | Group 2 | Difference in means | Standard error | T-value | Tukey-HSD corrected p-values | Cohen’s d |
| --- | --- | --- | --- | --- | --- | --- |
| FD-e | M | 0.05 | 0.09 | 0.58 | 0.83 | 0.12 |
| FD-e | UD | -0.24 | 0.08 | -2.96 | 0.0096 | -0.54 |
| M | UD | -0.29 | 0.08 | -3.59 | 0.0012 | -0.61 |

**Supplementary Table 8.** Summary table of post-hoc testing in Dataset 1b.

##### Dataset 2 – d’

We performed one-way ANOVA with the first condition as the between-subject factor and d’ as the dependent variable that didn’t show a significant effect of the first condition (F(2, 149)=0.25; p=0.78; MSE = 0.06; $\eta_{p}^{2}$= 0.0034). Subsequently, we performed a pairwise Tukey-HSD post-hoc test and we summarise the results in SI Table 9 below.

| Group 1 | Group 2 | Difference in means | Standard error | T-value | Tukey-HSD corrected p-values | Cohen’s d |
| --- | --- | --- | --- | --- | --- | --- |
| FD-e | M | -0.01 | 0.09 | -0.12 | 0.99 | -0.02 |
| FD-e | UD | -0.07 | 0.10 | -0.68 | 0.78 | -0.13 |
| M | UD | -0.06 | 0.10 | -0.57 | 0.84 | -0.11 |

**Supplementary Table 9.** Summary table of post-hoc testing in Dataset 2.

##### Dataset 3 – d’

We performed one-way ANOVA with the first condition as the between-subject factor and d’ as the dependent variable that showed a significant effect of the first condition (F(2, 259)=8.94; p=0.0002; MSE=1.86; $\eta_{p}^{2}$= 0.065). Subsequently, we performed a pairwise Tukey-HSD post-hoc test and we summarise the results in SI Table 10 below.

| Group 1 | Group 2 | Difference in means | Standard error | T-value | Tukey-HSD corrected p-values | Cohen’s d |
| --- | --- | --- | --- | --- | --- | --- |
| FD-e | M | 0.06 | 0.07 | 0.82 | 0.69 | 0.13 |
| FD-e | UD | -0.22 | 0.07 | -3.15 | 0.005 | -0.48 |
| M | UD | -0.28 | 0.07 | -4.02 | 0.0002 | -0.59 |

**Supplementary Table 10.** Summary table of post-hoc testing in Dataset 3.

##### Concatenated datasets 1a, 2, 3 – d’

We performed one-way ANOVA with the first condition as the between-subject factor and d’ as the dependent variable that showed a significant effect of the first condition (F(2, 593)=12.52; p=4.7$\cdot$10^-6^; MSE=2.71; $\eta_{p}^{2}$= 0.04). Subsequently, we performed a pairwise Tukey-HSD post-hoc test and we summarise the results in SI Table 11 below.

| Group 1 | Group 2 | Difference in means | Standard error | T-value | Tukey-HSD corrected p-values | Cohen’s d |
| --- | --- | --- | --- | --- | --- | --- |
| FD-e | M | 0.03 | 0.05 | 0.73 | 0.75 | 0.08 |
| FD-e | UD | -0.18 | 0.05 | -3.88 | 0.0003 | -0.38 |
| M | UD | -0.22 | 0.05 | -4.66 | 1.2$\cdot$10^-5^ | -0.45 |

**Supplementary Table 11.** Summary table of post-hoc testing in Concatenated datasets 1b, 2, 3.

#### Stimuli information

| **Database** | **Count** | **Proportion** |
| --- | --- | --- |
| CFD^107^ | 559 | 0.386 |
| BFD-BOGAZICI^104^ | 218 | 0.150 |
| Oslo^110^ | 147 | 0.101 |
| CFD India^109^ | 130 | 0.090 |
| London^58^ | 97 | 0.067 |
| 3DSK^103^ | 96 | 0.066 |
| RADIATE^111^ | 91 | 0.063 |
| CFD MR^108^ | 66 | 0.046 |
| PICS^134^ | 26 | 0.018 |
| KDEF^105^ | 19 | 0.013 |

**Supplementary Table 12.** Database representation. Count and percentage of images taken from the different databases.

| **Race** | **Count** | **Proportion** |
| --- | --- | --- |
| White | 521 | 0.3596 |
| Black | 242 | 0.1670 |
| Turkish | 216 | 0.1491 |
| Indian | 130 | 0.0897 |
| Latino | 105 | 0.0725 |
| Asian American | 101 | 0.0697 |
| Multiracial American | 66 | 0.0455 |
| Asian | 21 | 0.0145 |
| Hispanic | 18 | 0.0124 |
| West Asian | 10 | 0.0069 |
| East Asian | 8 | 0.0055 |
| Iranian | 8 | 0.0055 |
| Other | 3 | 0.0021 |

**Supplementary Table 13.** Race representation of face stimuli as reported in the databases.

### SI Figures

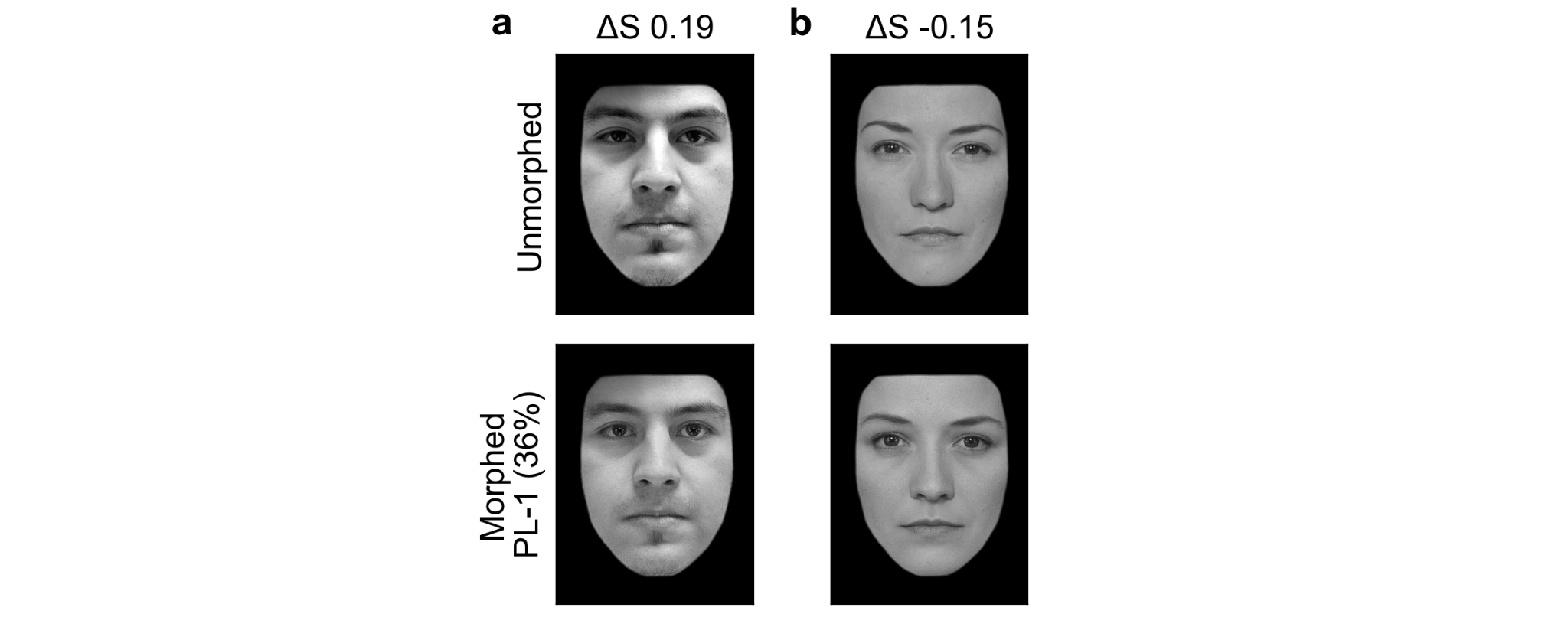

**Supplementary Figure 1.** Examples of stimuli with high positive (**a**, left column) and high negative (**b**, right column) difference in entropy between the Unmorphed image (top row) and the morphed image, in this case the most morphed face, 36% (bottom row). Image in **a** originates from the RADIATE^111^ database while image in **b** is taken from the London database^58^. Both are used in accordance with copyright.

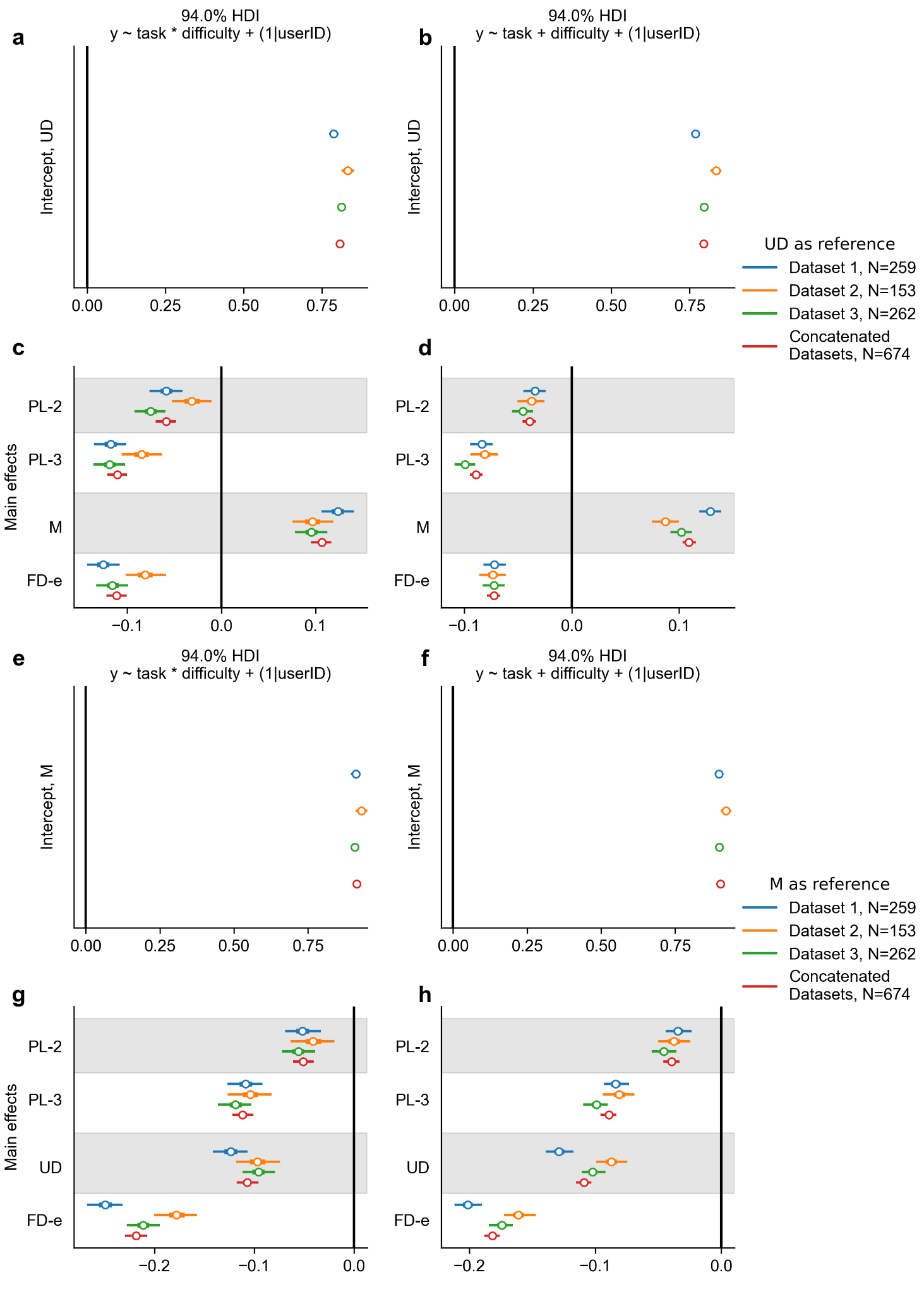

**Supplementary Figure 2.** Posterior distributions of main effects (memory condition and perceptual difficulty) across all three datasets as well as their concatenation show a significant effect of memory and perceptual difficulty levels (none of the distributions contains 0 in their 94% HDI in any of the datasets). Panels **a-d** show results for UD set as a reference level while panels **e-h** show the same plots with M set as a reference level. Top row shows the intercept value that represents UD level 1 and is significantly different from 0 (black line). Bottom row then shows two levels of each main effect – difficulty levels 2 and 3; memory condition of M and FD-e. Left column shows a model with interaction between memory and perception, the right column without. Both show comparable results with significant main effects. Black line passing through zero represents the reference level of UD level 1. Shown are two models – with interaction (panels **a, c**) and without (panels **b, d**), the equation for each is written in the title.

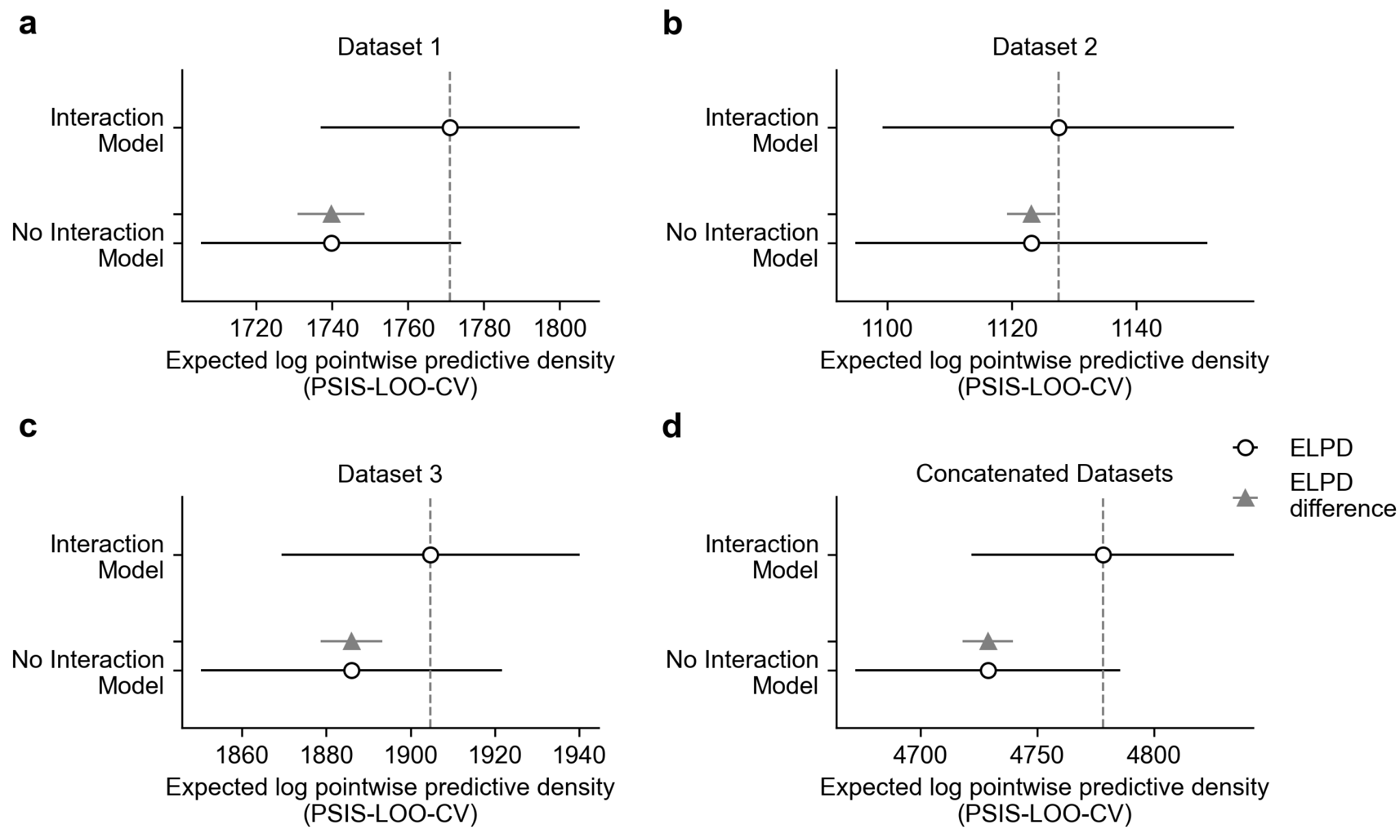

**Supplementary Figure 3.** Model comparison of two models, one with and one without the interaction term, shows that the model with included interaction performs better than the model without in three tested datasets (Dataset 1 in **a**, Dataset 3 in **c**, concatenated datasets in **d**) and no significant difference between these two models in Dataset 2 in panel **b**. ELPD was estimated by Pareto smoothed importance sampling leave-one-out cross-validation (PSIS-LOO-CV).

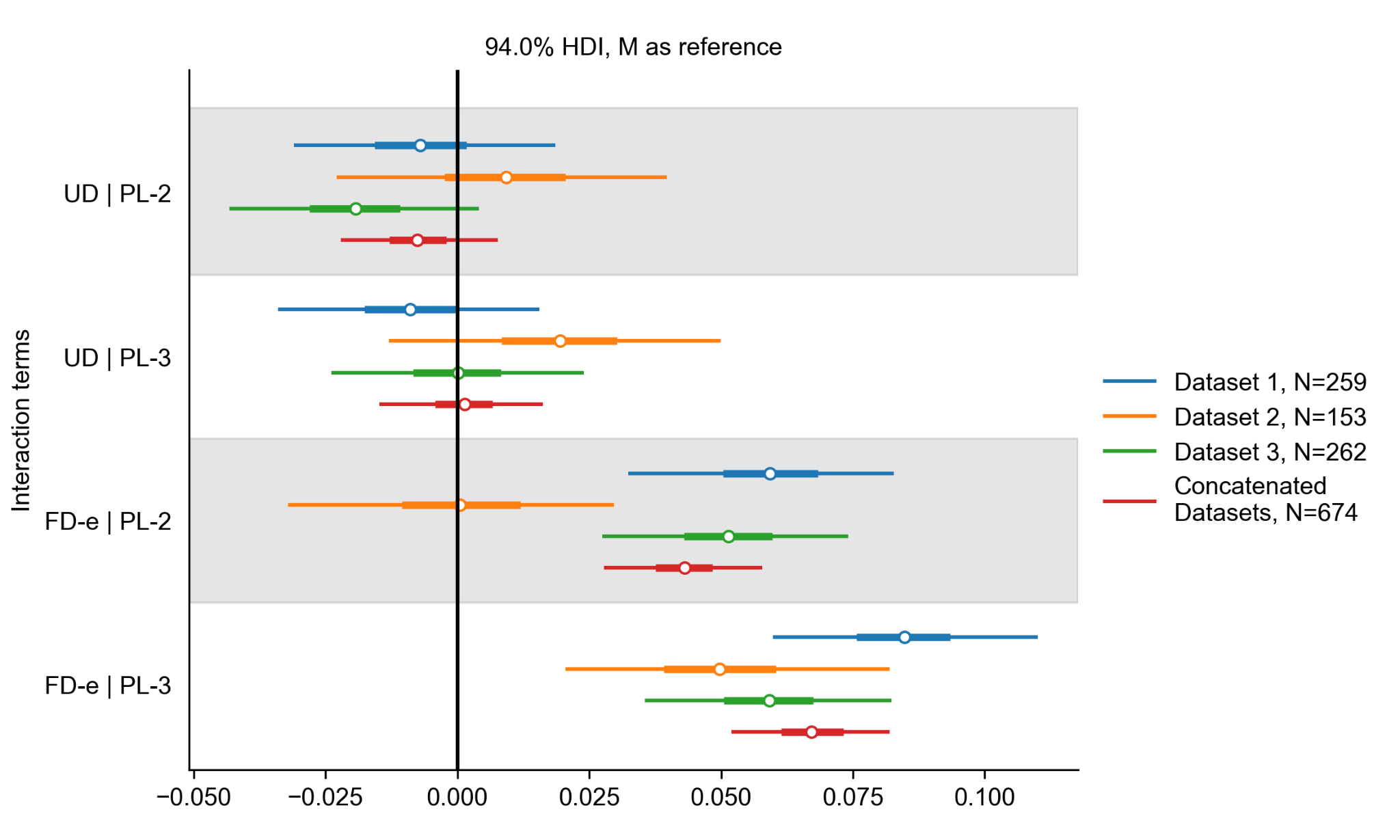

**Supplementary Figure 4.** Posterior distributions of interaction terms across the three datasets and the concatenated dataset show that there is no significant interaction between UD and M and the perceptual difficulty levels (both UD difficulty 2 and 3 and a significant interaction between M and FD-e and perceptual difficulty except for Dataset 2 perceptual level 2. Condition M set as reference.

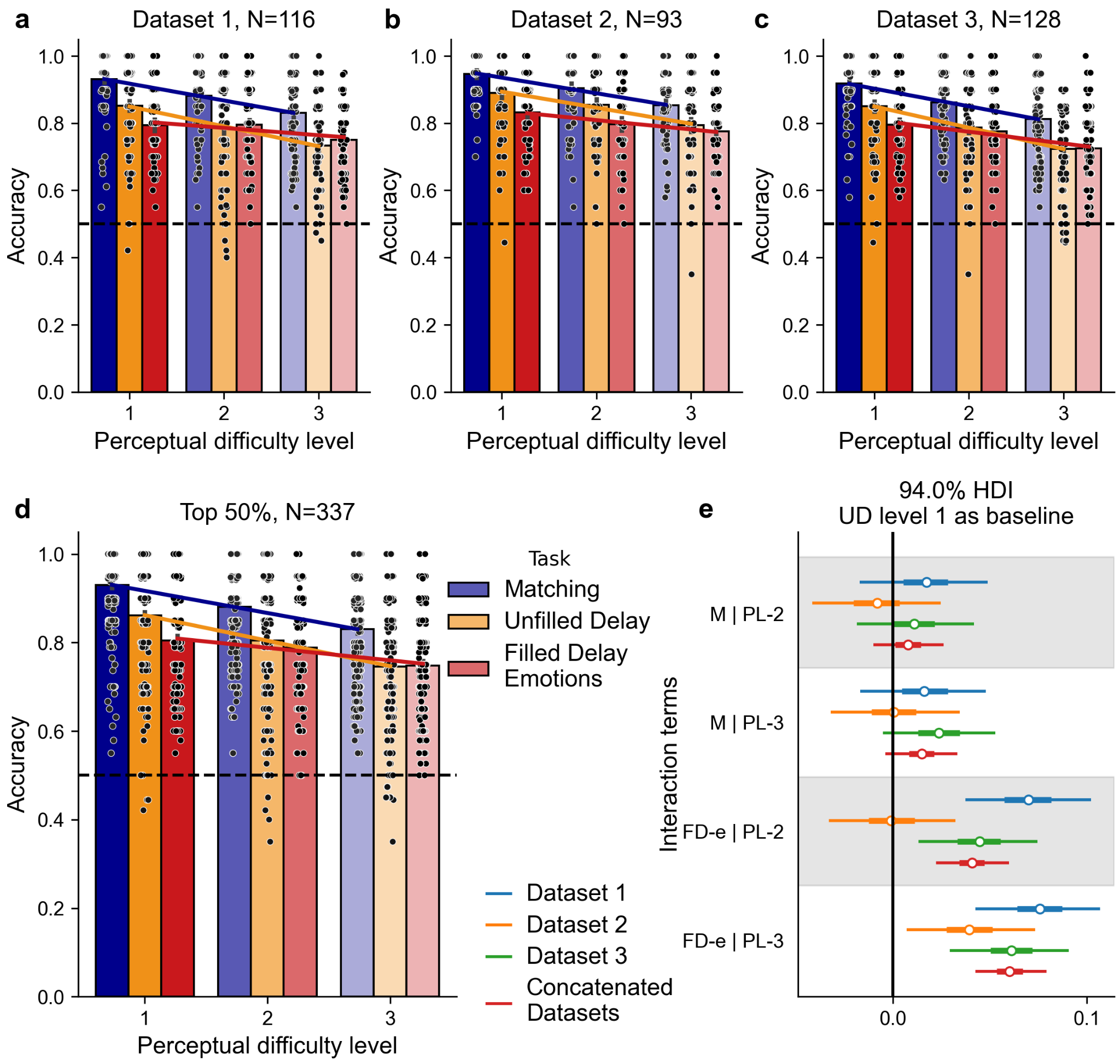

**Supplementary Figure 5.** To ensure that the results in Figure 2 are not a result of floor or restricted range effects, we matched the accuracy on the FD-e level 1 to UD level 1, by selecting the top 50% of participants on the FD-e condition and reran the analysis. Results on this subset show independence of Matching (M) and Unfilled Delay (UD) across perceptual difficulty levels (PLs) but not of the Filled Delay (FD-e) memory condition and perceptual difficulty. Each dot represents a single participant's score that is computed as the mean of 20 trials. As in Figure 2, perceptual difficulty levels are as follows: level 1 (PL-1, easiest, 36% morph), level 2 (PL-2, 30% morph), and level 3 (PL-3, hardest, 24% morph). **a-c** three independent datasets, see Methods for details. **d.** Concatenated datasets 1, 2, and 3. Connecting lines serve as a visualisation of interactions, where parallel lines reflect independence of memory and perception (M and UD), while non-parallel slopes across memory conditions reflect interaction between the processes (FD-e and M, FD-e and UD). **e.** Posterior distributions of interaction terms across all three datasets as well as their concatenation show no significant interaction between UD level 1 and M at either of the perceptual difficulty levels 2 or 3 (M | PL-2 and M | PL-3, respectively; posterior distributions contain 0 in their 94% HDI) and that there is a significant interaction between UD and FD-e and perceptual difficulty except for Dataset 2 on level 2. UD perceptual difficulty level 1 is set as baseline.

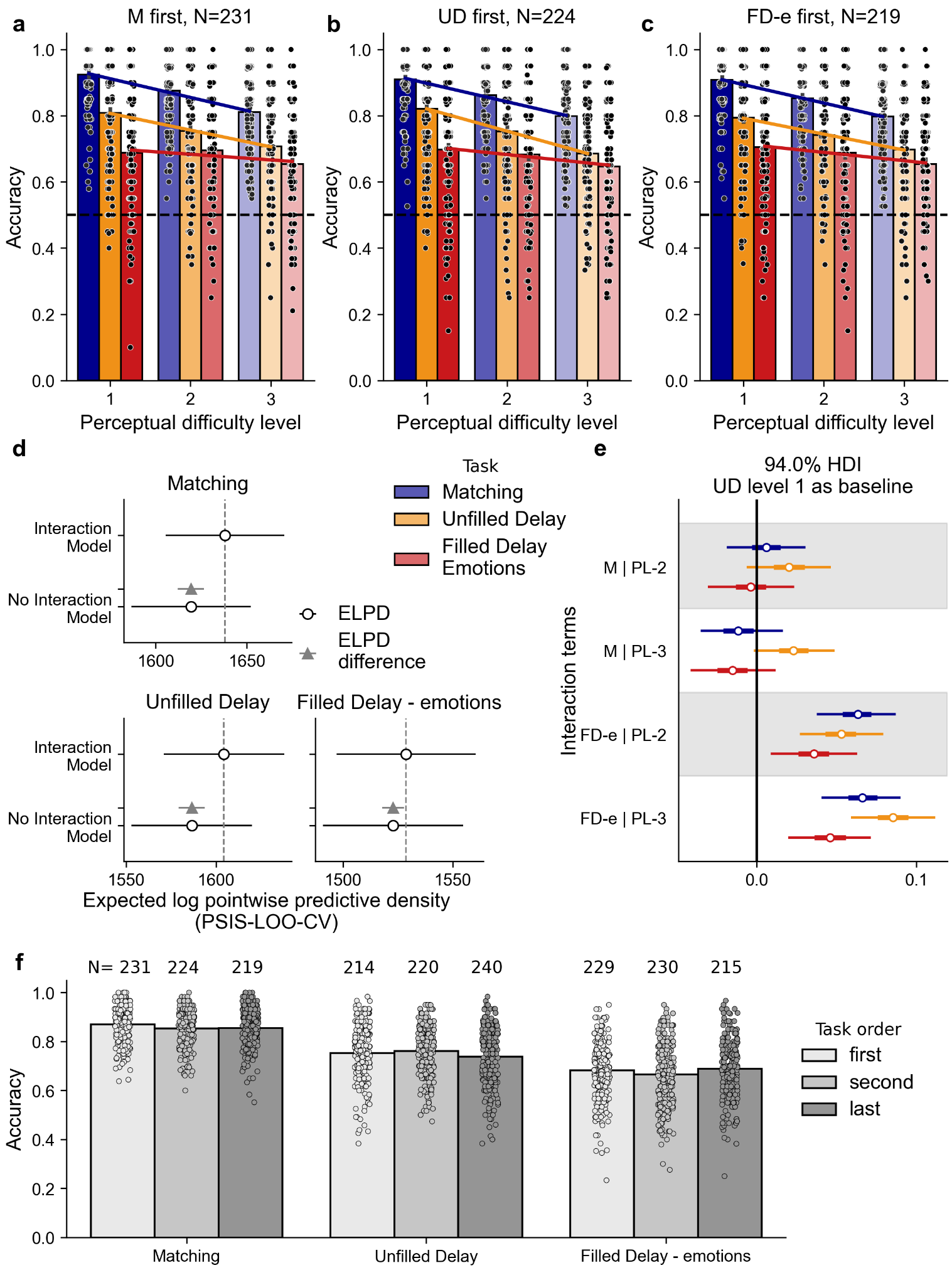

**Supplementary Figure 6.** Robustness of main results across condition order and participant subsets. (a–c) Results for three independent subsets of the concatenated dataset, grouped by participants who began with the M condition (**a**), the UD condition (**b**), or the FD-e condition (**c**). **d.** Model comparison showing the model with the interaction term consistently outperforms the model without, across all three sub-datasets. **e.** Posterior distributions of interaction terms across the three sub-datasets show no significant interaction between UD level 1 and M at either of the perceptual difficulty levels 2 or 3 (M | PL-2 and M | PL-3, respectively; posterior distributions contain 0 in their 94% HDI) and that there is a significant interaction between UD and FD-e and perceptual difficulty. UD perceptual difficulty level 1 is set as baseline. **f.** Accuracy by condition order (first, second, last) for M, UD, and FD-e conditions shows there is no consistent pattern (increase or decrease of accuracy with time) suggesting there is no learning present, see Supplementary Table 7 for stats. N for each group is indicated above the bars.

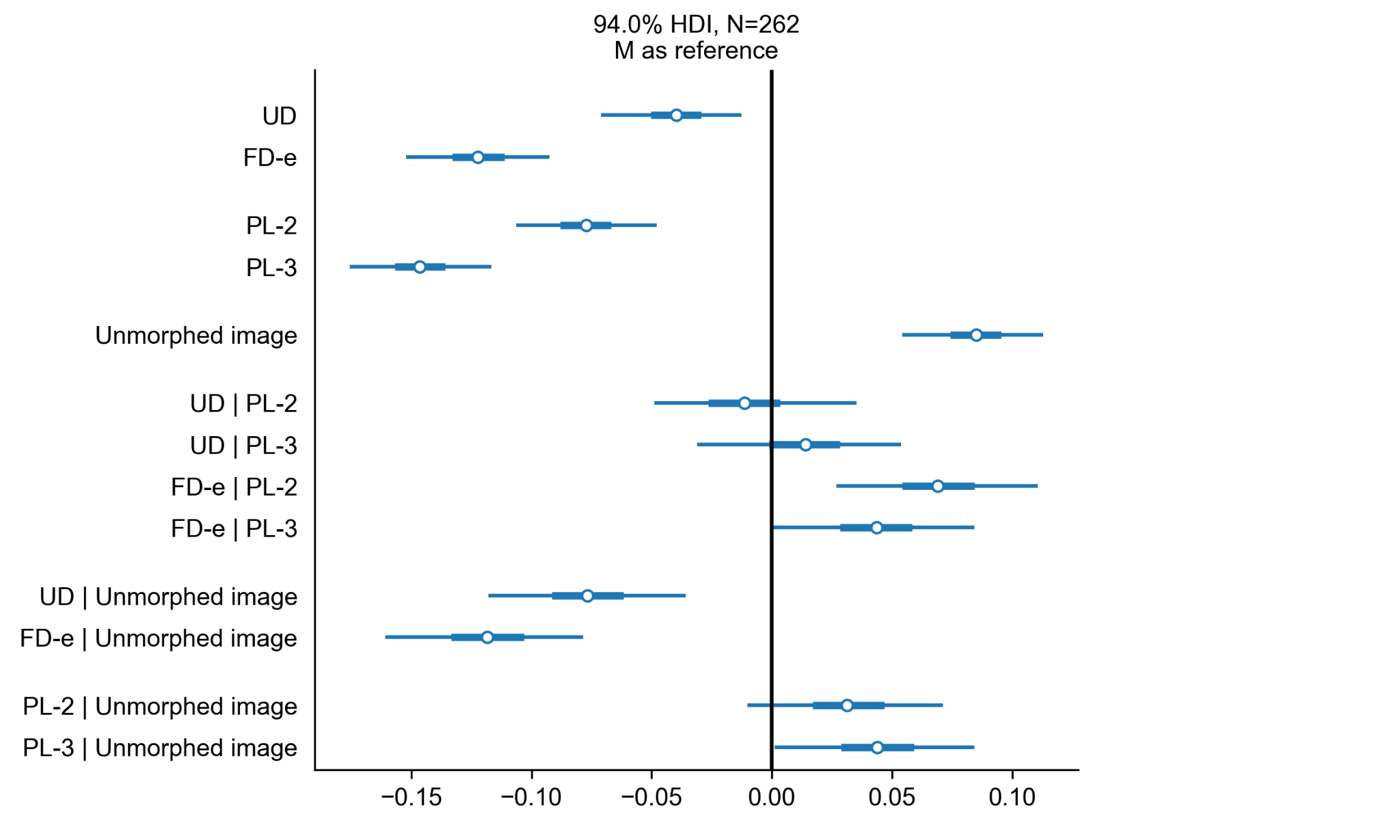
**Supplementary Figure 7.** The model shows the main effects of memory condition (M/UD, FD-e) and perceptual difficulty (levels 2, 3). When M is set as a reference condition, we see a significant effect of the cue stimuli and a significant interaction between the cue stimulus and both memory levels (UD, FD-e) and perceptual difficulty on level 3. Consistently with Figure 3e, there is no interaction between UD and M and perceptual levels, and there is a significant interaction between M and FD-e and both perceptual levels.

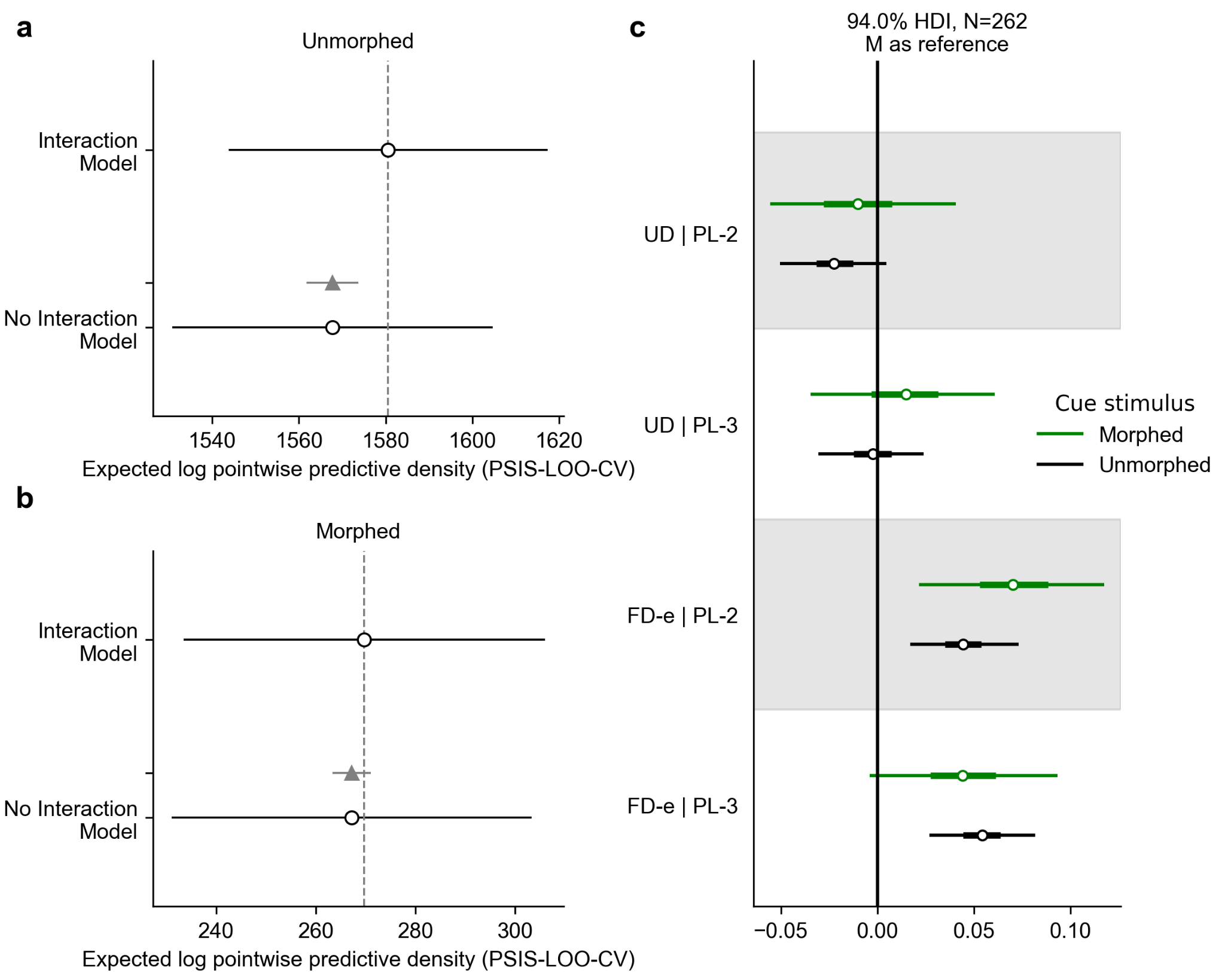

**Supplementary Figure 8. a.** Model comparison of two models, one with and one without the interaction term, shows no significant difference between these two models in Dataset 3 that looks separately at cue stimulus. **b.** There is a small difference between the two models when the cue stimulus is unmorphed (original) and no significant difference when the cue stimulus is morphed. **c.** Posterior distributions of interaction terms show no significant interaction as the 94% HDI contains zero in both datasets between UD and M with M set as reference level. The FD-e condition behaves differently in the dataset with morphed cue stimuli as there is no significant interaction between level 3 but there is in level 2. The findings align with those in Figure 3f.

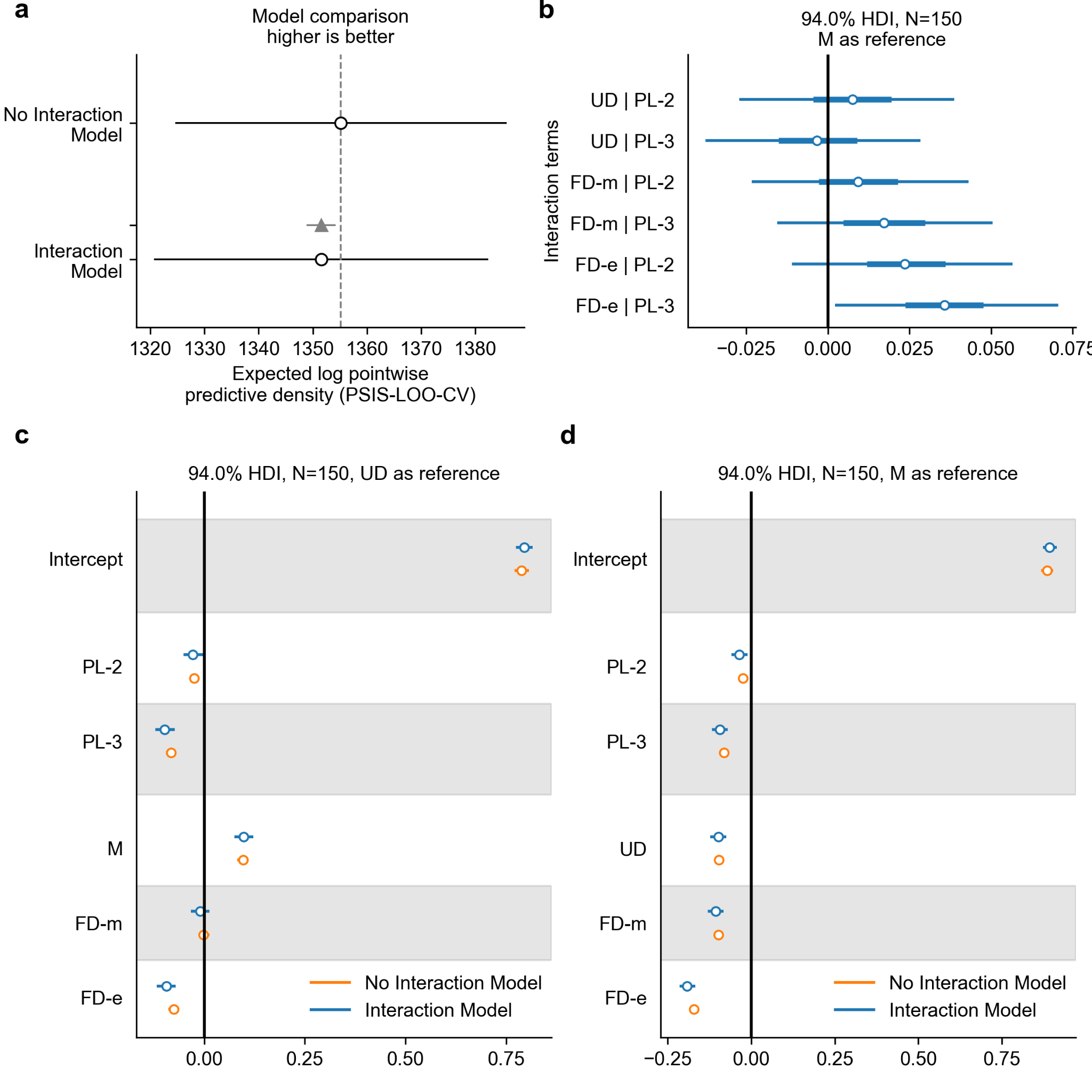

**Supplementary Figure 9. a.** Model comparison of two models, one with and one without the interaction term, shows that model without the interaction performs slightly better than the model with interaction in Dataset 4 (Math interference). ELPD was estimated by Pareto smoothed importance sampling leave-one-out cross-validation (PSIS-LOO-CV). **b.** Posterior distributions of interaction terms show no significant interaction as the 94% HDI of all of them contain zero (black line) except for FD-e level 3. Interaction terms are defined as condition:difficulty level. Zero corresponds to M level 1. **c.** Posterior distributions for all memory conditions and perceptual difficulties show no difference between FD-m and UD conditions as the 94% HDI is centred at zero (black line). This panel also shows the main effect of memory difficulty (M and FD-e being significantly different from zero) as well as perceptual difficulty (level 3 being significantly different from 0). **d.** Same as **c.** but with M set as a reference.

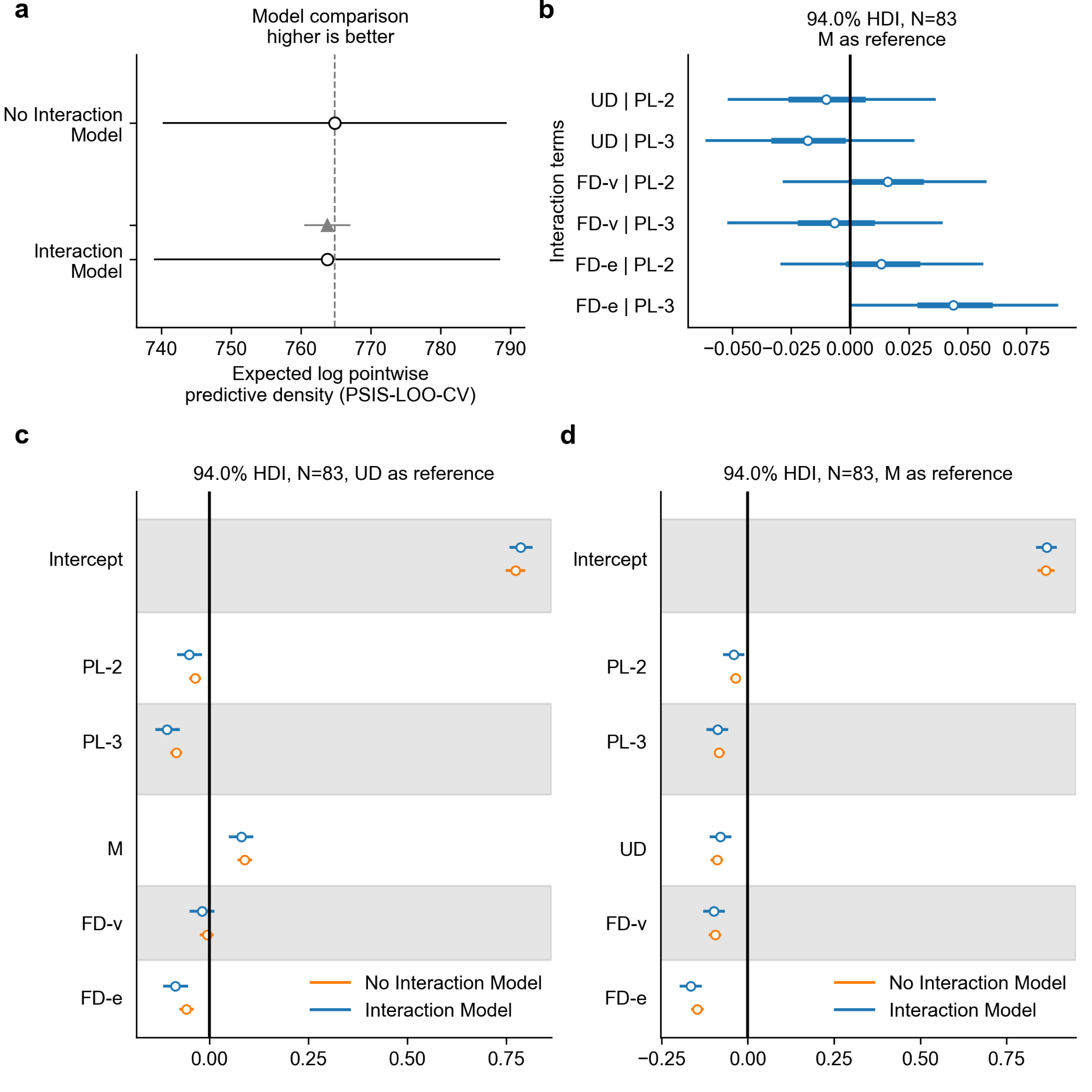

**Supplementary Figure 10. a.** Model comparison of two models, one with and one without the interaction term, shows that model without the interaction performs no better than the model with interaction in Dataset 5 (Visual interference with food plate distractor). ELPD was estimated by Pareto smoothed importance sampling leave-one-out cross-validation (PSIS-LOO-CV). **b.** Posterior distributions of interaction terms show no significant interaction as the 94% HDI of all of them contain zero (black line) except for FD-e level 3. Interaction terms are defined as condition:difficulty level. Zero corresponds to M level 1. **c.** Posterior distributions for all memory conditions and perceptual difficulties show no difference between FD-v and UD conditions as the 94% HDI is centred at zero (black line). This panel also shows the main effect of memory difficulty (M and FD-e being significantly different from zero) as well as perceptual difficulty (level 3 being significantly different from 0). **d.** Same as **c.** but with M set as a reference.

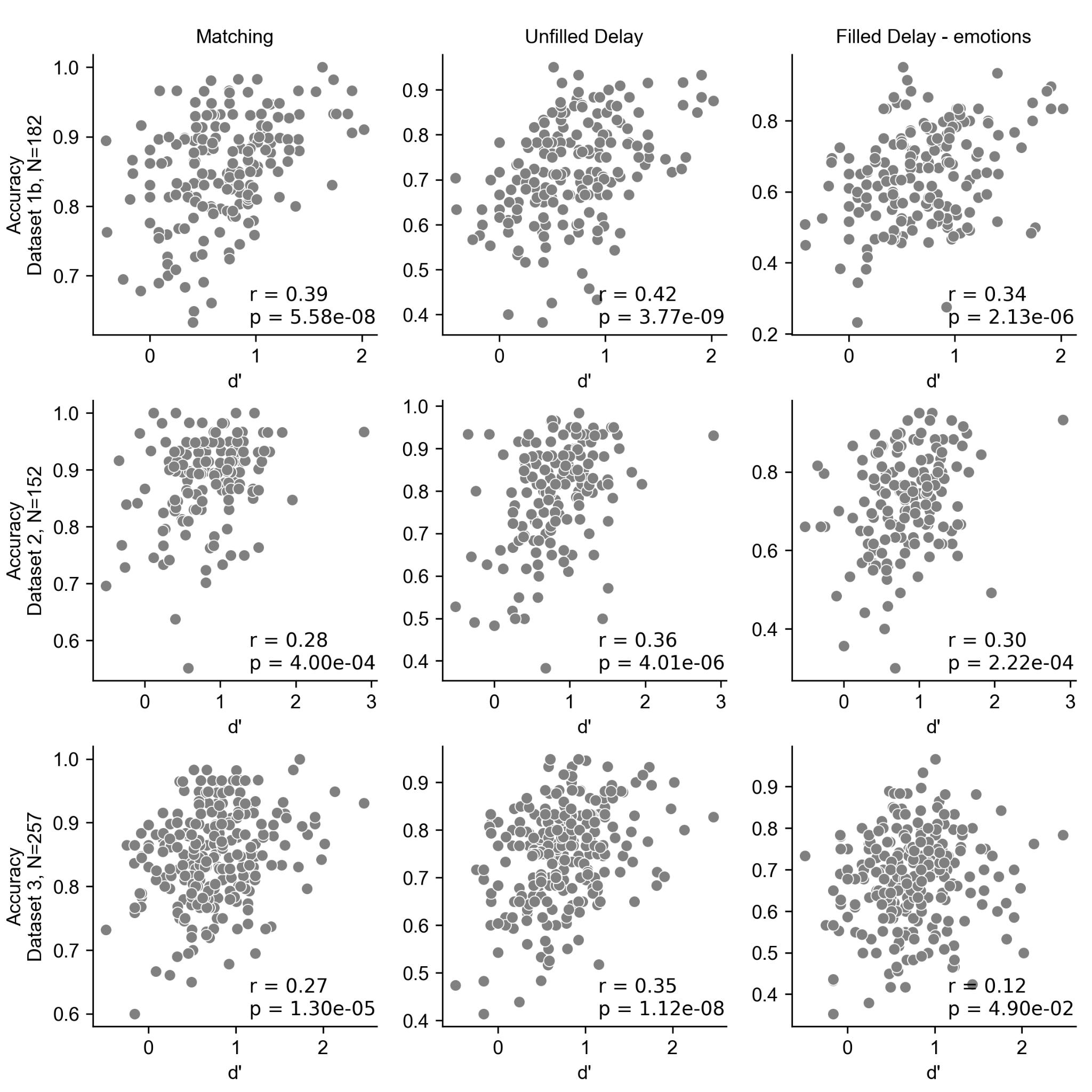

**Supplementary Figure 11.** Correlation between d’ on the Long-term recognition memory task and accuracy in the memory conditions (M, UD, FD-e, columns) across the three datasets (rows).

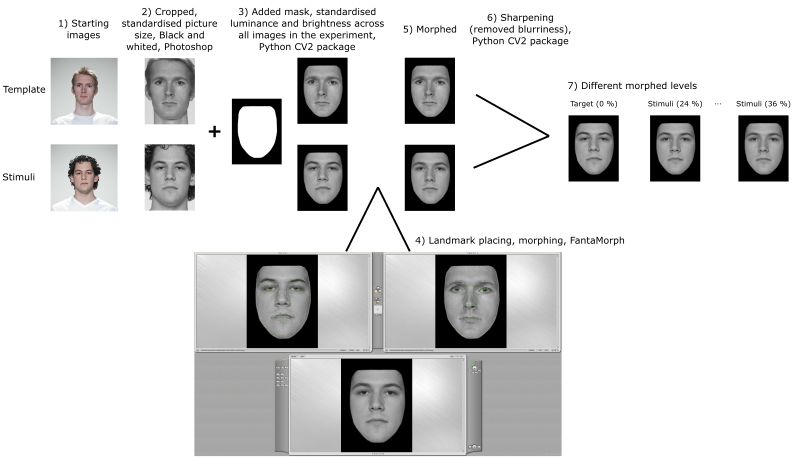

**Supplementary Figure 12.** Stimuli creation. All pictures (1) were first manually cropped, resized, converted to black and white and all significant birthmarks were removed using Adobe Photoshop (2). In the next step (3), images were standardised for luminance and brightness and a mask was added over them using python and cv2 package. Subsequently, 99 landmarks were placed on the face outlining the face, the eyebrows, the eyes, the iris, the mouth, and the nose. This was done manually in FantaMorph (4). One female and one male face were chosen as template images (top row) and all faces were morphed towards these two template faces (5). We used 24%, 30%, and 36% morphs. After morphing, stimuli were sharpened twice using the cv2 package in python (6) which created a set of 4 images – original (unmorphed) target face and three morphed faces (7). Shown images were taken from the London database^58^ and are presented in accordance with copyright.

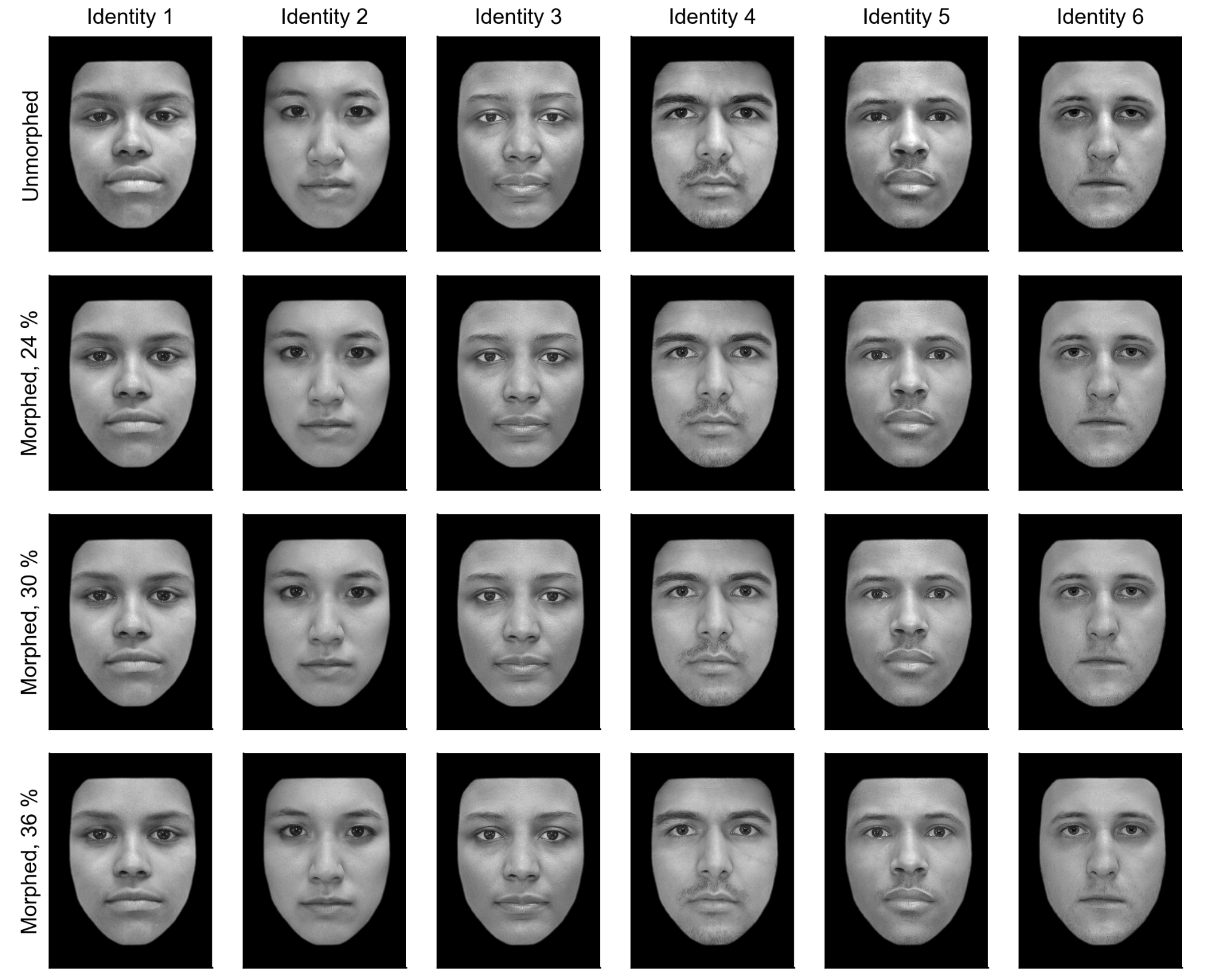

**Supplementary Figure 13.** Gallery of example morphed stimuli used in the experiment. All images were taken from the RADIATE^111^ database of faces and are used in accordance with copyright. Each column displays one identity – identities 1-3 are female faces, identities 4-6 are male faces. Each row represents different levels of morphing with the first being unmorphed (original image), second corresponds to 24% (PL-3), third to 30% (PL-2) and fourth to 36% (PL-1). The race according to the database of the identities goes from left to right: Hispanic, Asian, Black, Asian, Black, White.

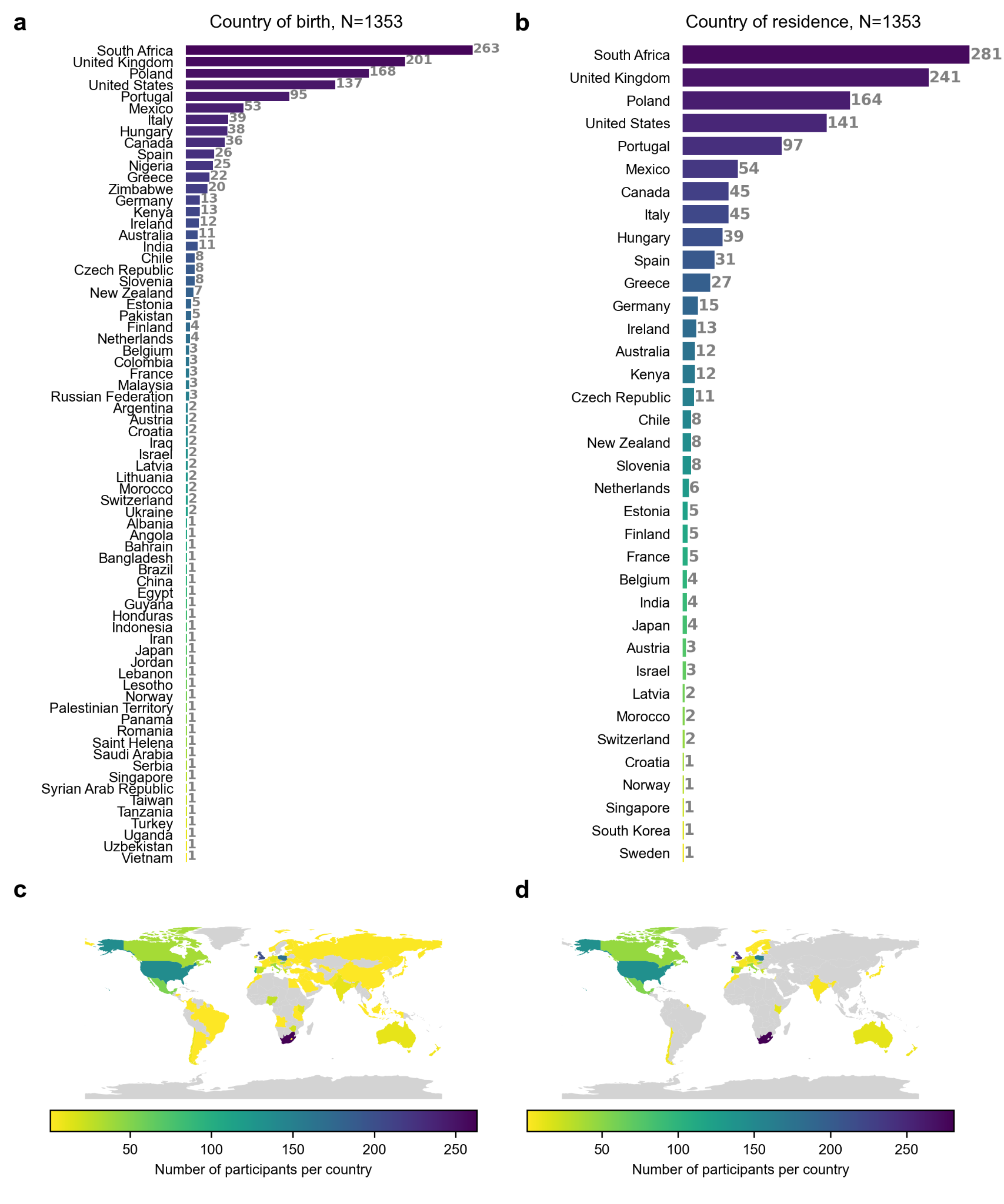

**Supplementary Figure 14.** Demographics of the participants involved in the study. Panels **a** and **c** display the countries of birth, while panels **b** and **d** show the participants' current countries of residence. This figure is intended as a tribute and an expression of gratitude to all those who contributed to the study, although not all participants are represented here.
